# Dimension-agnostic and granularity-based spatially variable gene identification

**DOI:** 10.1101/2023.03.21.533713

**Authors:** Juexin Wang, Jinpu Li, Skyler T Kramer, Li Su, Yuzhou Chang, Chunhui Xu, Qin Ma, Dong Xu

## Abstract

Identifying spatially variable genes (SVGs) is critical in linking molecular cell functions with tissue phenotypes. Spatially resolved transcriptomics captures cellular-level gene expression with corresponding spatial coordinates in two or three dimensions and can be used to infer SVGs effectively. However, current computational methods may not achieve reliable results and often cannot handle three-dimensional spatial transcriptomic data. Here we introduce BSP (big-small patch), a spatial granularity-guided and non-parametric model to identify SVGs from two or three-dimensional spatial transcriptomics data in a fast and robust manner. This new method has been extensively tested in simulations, demonstrating superior accuracy, robustness, and high efficiency. BSP is further validated by substantiated biological discoveries in cancer, neural science, rheumatoid arthritis, and kidney studies with various types of spatial transcriptomics technologies.

## Background

Spatially resolved transcriptomics (SRT) have been rapidly developed and widely used in biological and biomedical research over the past decade^1-3^. Single-molecule fluorescence *in situ* hybridization (smFISH) (e.g., MERFISH and SeqFISH+) and sequencing-based approaches (e.g., 10X Visium)^2^ are popular SRT technologies on sliced two-dimensional (2D) samples. A shift has recently occurred towards retaining three-dimensional (3D) positional anatomy at cellular resolution. Wang *et al*. developed STARmap, which combined an efficient sequencing approach with hydrogel-tissue chemistry for 3D intact tissue RNA sequencing^4^, with a throughput of up to 1,000 genes or 10,000 genes^5^. Vickovic *et al*. developed protocols on consecutive sections to get 3D spatial profiling of rheumatoid arthritis (RA) synovia^6^. Existing works on 3D imaging data construction and 3D imaging data have significant advantages over 2D data for accurate quantitative interpretation^7^. In contrast to the 2D spatial transcriptomics approach, which depends on sampling strategy (e.g., coronal or sagittal) on sliced samples, the 3D spatial transcriptomics provides a more comprehensive and faithful representation of intact organ structures and functions. It overcomes the inherent 2D bias and enables the visualization of gene expression in relation to the tissue architecture in three dimensions. Such 3D views provide new opportunities in the identification of cell types and states, discovery of new biomarkers, and drug design^8^. However, most of the current analytic methods are developed and validated on 2D SRT data, and cannot be directly applied to diverse types of 3D analyses^9^.

Spatially variable genes (SVGs) are biologically significant as they exhibit variations in expression levels across different regions or cell types within a tissue, indicating their involvement in specific biological processes or functions unique to those regions or cell types. Hence, the inference of SVGs can help researchers gain a deeper understanding of how different cell types and genes contribute to the overall structures and functions of tissues in normal or disease states^10^. Additionally, SVGs can be used as molecular markers to track developmental or disease-related changes in the spatial distribution of specific cell types. Identification of SVGs also facilitates the dissection of biological relationships between spatial organization and molecular cell function, providing critical information for biologists and pathologists. For example, in the mouse olfactory bulb, Stahl *et al*. discovered functional regions in the mouse brain by identifying SVGs^11^, while Maynard *et al*. discovered laminar and nonlaminar genes in the human dorsolateral prefrontal cortex^12^. The existing SRT technologies encode the key clues of SVGs, whose expressions rely on the spatial locations of cells^13,14^, and identifying SVGs from tens of thousands of genes is often a critical first step in analyzing spatial transcriptome data.

Compared to traditional RNA-seq and scRNA-seq analyses that identify differentially expressed (DE) genes, SVGs in SRT data incorporate gene expression information and corresponding spatial context in geometric coordinates. In 2D SRT studies, SpatialDE^13^ and Trendsceek^14^ are the first two computational methods for identifying SVGs: SpatialDE utilizes Gaussian process regression to quantify the spatial variance of expression for each gene, while Trendsceek selects SVGs by testing the dependence between the expression and the spatial location for each gene using a permutation process. Afterward, a generalized linear spatial model with Gaussian/periodic kernels, SPARK^15^, was proposed to capture the spatial patterns and filter SVGs using the combined *p*-values from each kernel. A simplified version, SPARK-X^16^, was later introduced to reduce computational time and memory usage. In addition to statistical methods, MERINGUE^9^ applied a Voronoi tessellation method to build an adjacency matrix and calculate classical Moran’s I score^17^ for each gene based on the constructed adjacent matrix to infer SVGs. SpaGCN^18^ first identifies the spatial domain with graph neural networks and then employs a statistical test to identify SVGs based on the context of the inferred spatial domains.

Although some preliminary analysis^4,6^ has been conducted on emerging 3D SRT data, significant challenges remain in identifying SVGs in the dimension-agnostic SRT data^9,16^. The limited spatial information captured by 2D tissue slices may result in incomplete and biased representations of spatial characteristics, potentially leading to inaccurate biological conclusions^8,19^. Additionally, the existing SVG identification methods require user-defined parameters that can vary across samples and lead to disparate findings that are difficult to justify without prior knowledge of the samples. Hence, a non-parameter method with adequate power is preferred even for 2D data in practical usage. Based on our preliminary analysis, the expression distribution of SVGs tends to exhibit a consistent and specific pattern invariant across different spatial resolutions and views, whereas the expression distribution of non-SVGs has a random pattern with varying characteristics across different views and resolutions. These distributions can be effectively captured by granularity, a concept relatively underexplored in spatial transcriptomics studies. Granularity refers to the extent or hierarchical level to which a material or system comprises distinguishable pieces^20^. We propose that the concept of granularity can be leveraged to identify SVGs in a dimension-agnostic geometric manner. With appropriate quantitative measures, granularity-based criteria can distinguish between spatial organizations with biological significance and those with random patterns.

Here we introduce BSP (big-small patch), a spatial granularity-guided and non-parametric model that enables efficient and robust identification of SVGs from two/three-dimensional SRT data. For each spot in the data, BSP selects a set of neighboring spots within a certain distance to capture the regional means with different granularities. The variances of the expression mean across all spots are then calculated under different scales, and genes with high ratios are identified as the SVGs. One of the unique features of BSP is that it does not make any assumption regarding the distribution of the gene expression levels or the spatial pattern of the spots. The model is robust to fluorescence *in situ* hybridization (MERFISH, seqFISH+, and STARMap) and sequencing-based (10X Visium and slide-seq) SRT without requiring pre-defined or well-tuned parameters. Compared with existing methods, BSP outperforms other methods for 3D data, and delivers comparable power and accuracy as current methods for 2D data with a significantly reduced computational cost. In addition, the BSP algorithm is easily implementable, making it versatile and easily integrated into various applications. In our experiments on kidney SRT data and 3D RA synovia study, BSP identified several functional-related SVGs related to the disease mechanisms. In summary, BSP is an accurate, fast, robust, and parameter-free method for identifying SVGs in 2D and 3D SRT data.

## 2. Results

### 2.1 The big-small-patch algorithm

The proposed BSP algorithm is a granularity-guided approach for identifying SVGs in dimension-agnostic SRT data (**Figure 1**). BSP defines a patch for each spot in the SRT data, which includes all neighboring spots within a given radius centered on the spot (**Figure 1A**). A pair of patches is then defined, consisting of a small patch with a smaller radius and a large patch with a larger radius (**Figure 1B**). This paired big-small patch captures the ambient local expression characteristics in different granularities, delineating spatial patterns in various contexts. Subsequently, the transcriptomic expression variance of the local means is calculated across all pairs of patches, and the ratio between the variance with a big patch and the variance with a small patch is used as the statistic score for each gene. This statistic score can be used to quantify the conservation of SVGs’ spatial patterns in different granularities. The distribution of this statistical score is fitted with a beta distribution, and genes with statistical significance (*p* < 0.05) in the fitted beta distribution are defined as SVGs (**Figure 1C**).

**Figure 1.**
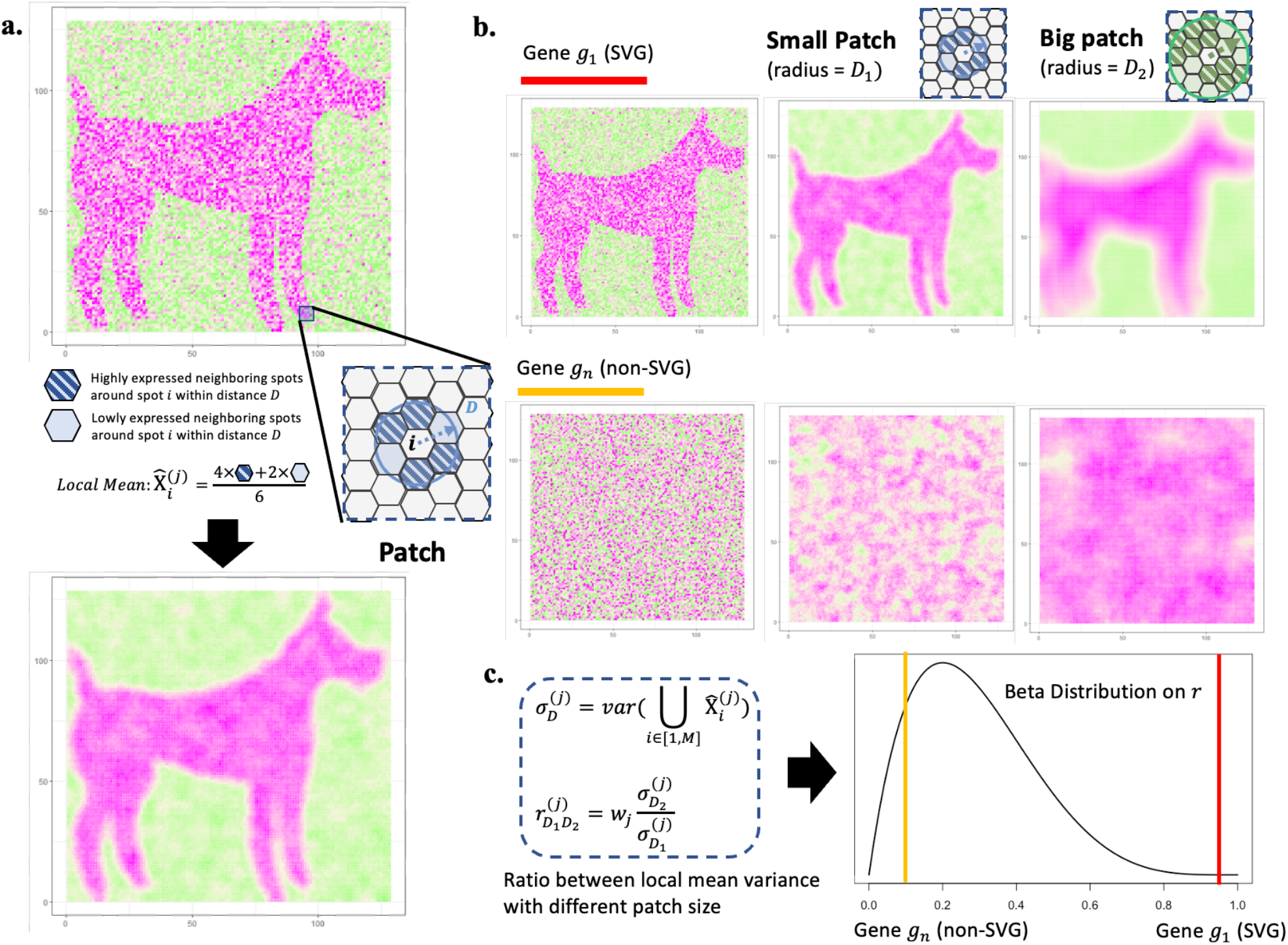
Scheme of BSP. **a)** Definition of the patch in BSP. For spot *i*, a patch is defined as the set of all its neighboring spots within distance *D*. Local Mean 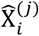 is defined as the average expression of gene *j* for all spots within the patch. **b)** Identification of SVGs with BSP. Gene *g*_1_ indicates an SVG (red), while *g*_*n*_ is a representative non-SVG gene (orange). Besides the original pattern as the left, we define a small patch of spots with a smaller radius *D*_1_ (blue) in the middle column, and a big batch of spots with a larger radius*D*_*2*_ (green) in the right column. **c)** The ratio 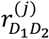 of the variance of the local mean between big batch 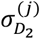 and the paired small batch 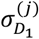 is chosen as the statistical value *r*. Gene *g*_%_ (red bar) is statistically significant compared with the background gene *g*_*n*_ (orange bar) on the fitted beta distribution of *r*.

### 2.2 BSP can accurately and efficiently identify SVGs in 2D simulations

To demonstrate the effectiveness of the BSP model in analyzing 2D transcriptomic data, we generated a set of simulations following the SPARK^15^ framework. We then compared the performances of the BSP model with a basic spatial statistic (Moran’s I^17^, which is also adopted by MERINGUE^9^) and other established techniques, including SpatialDE^13^, SPARK^15^, and SPARK-X^16^. To ensure a fair comparison of the model performances, we measured their statistical power based on the false discovery rate (FDR), considering the differences between the distribution of calibrated *p*-values from each method. We present three SVG patterns from previous works on the ST mouse olfactory analysis by SpatialDE and SPARK in **Figure 2A**. Details of the simulations are outlined in the Methods section.

**Figure 2:**
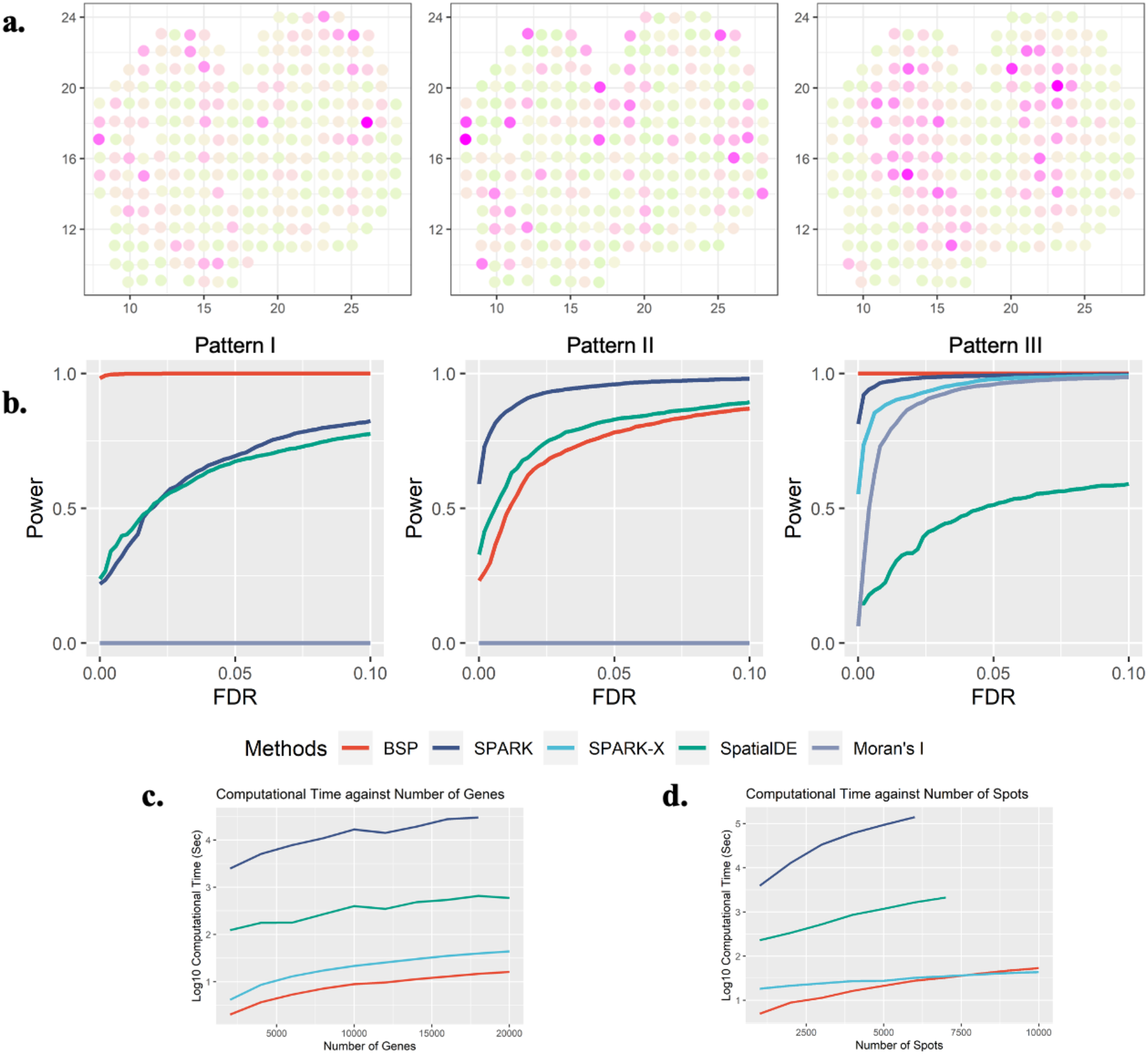
Power analysis for 2D simulation. **a)** Spatial expression patterns I-III (left to right) as defined in SpatialDE and SPARK. **b)** Power comparison among the different methods with moderate noise level (τ=0.5), and moderate signal strengths (3-folds) for the spatial expression patterns I, II, and III (left to right). All simulations were generated based on the mouse olfactory bulb data with 260 spots of cells. Each simulation replicate contains 1,000 SVGs and 9,000 non-SVGs. **c)** Computational time costs with an increasing number of genes and fixed 2,000 spots. The y-axis indicates the logarithmic of the running time in seconds. **d)** Computational time costs with an increasing number of spots and fixed 10,000 genes. The y-axis indicates the logarithmic of the running time in seconds. The time cost greater than 48 hours are not shown in the figure.

We evaluated the statistical power of different methods using simulations generated with various signal strengths and noise levels. Signal strengths were measured as the fold-change (*FC*) in cells’ expression levels between the pattern and non-pattern areas. To compare the statistical power, we examined different signal-to-noise ratios (*FC* = 3,4,5) with a moderate noise level (τ = 0.5 as defined in SPARK) in **Supplementary Figure 1**. Additionally, we compared the methods’ performance under different noise levels (τ = 0.2,0.5,0.8) with a moderate signal-to-noise ratio (*FC* = 4), as shown in **Supplementary Figure 2**. The BSP method consistently showed superior and stable power across a wide range of FDR cutoffs, signal strengths, and noise levels when analyzing the first and third spatial patterns (**Figure 2B**). Furthermore, BSP exhibited comparative power as SPARK and SpatialDE on the second pattern when the signal strength was moderate (3-fold) and the noise level τ equals 0.5. Compared to other existing approaches, BSP was more powerful, particularly on samples with low signal strengths or high noise levels.

We assessed the computational time and memory usage required for detecting SVGs on 2D data. Compared to existing approaches, BSP performs the SVG analysis with a feasible computational time and memory consumption on personal computers in most scenarios. Computational resource consumption was recorded on an Ubuntu 16.04.4 LTS workstation with Intel(R) Xeon(R) W-2125 CPU @ 4.00GHz and 32 GB memory in **Supplementary Table 1**. In analyzing a typical spatial transcriptomic sample with 2,000 spots, BSP was much faster than other existing methods, regardless of the number of genes (**Figure 2C**). Similarly, for a spatial transcriptomics sample with 10,000 genes, BSP had the lowest computational time among all methods, despite the number of spots (**Figure 2D**). Although not as low as SPARK-X, the corresponding memory usage was also low, as shown in **Supplementary Figure 3**.

**Figure 3.**
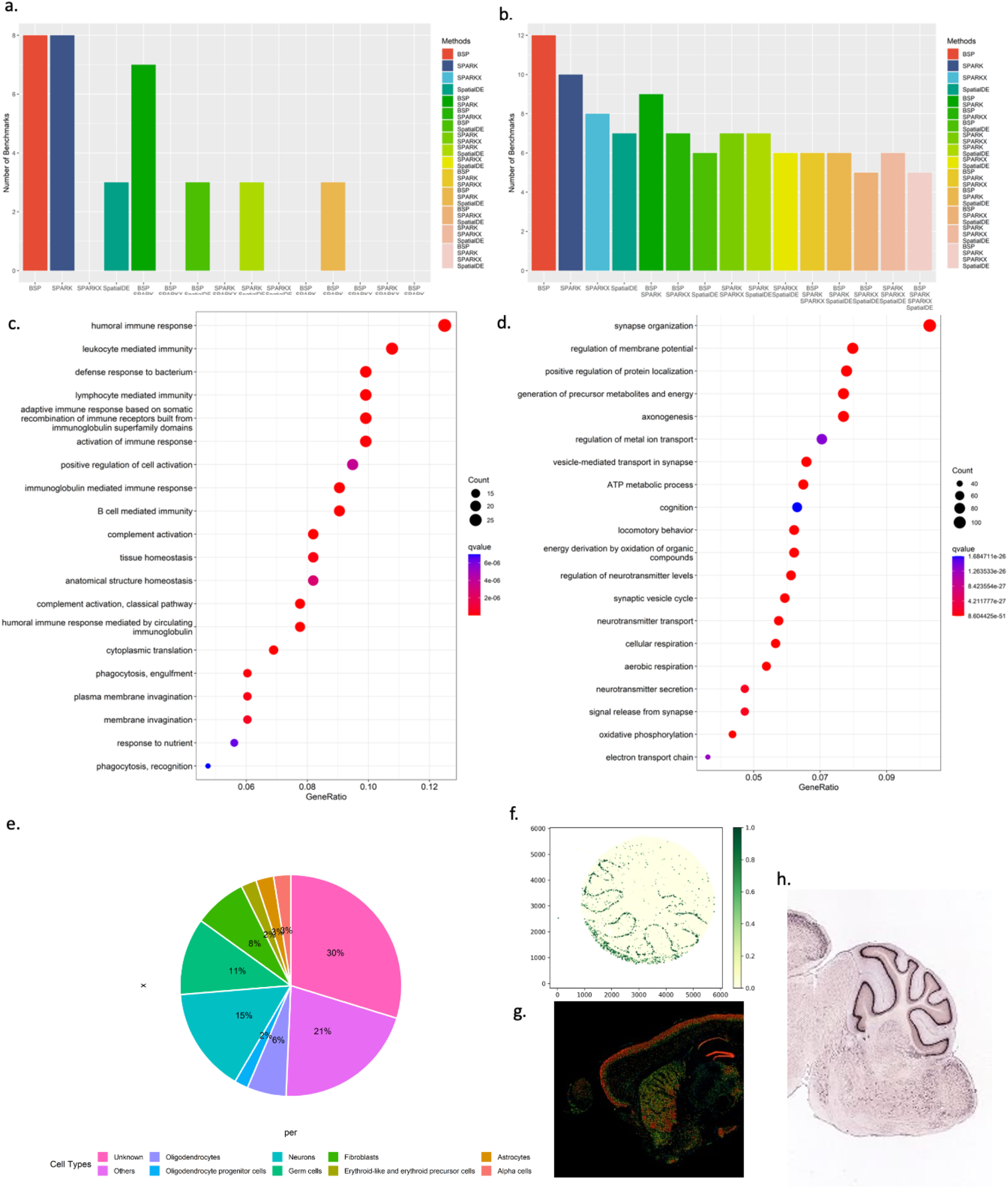
SVGs identified by BSP in biological analysis. **a)** Number of marker genes identified with different methods in mouse olfactory bulb study (the original study identified 10 marker genes). **b)** Number of marker genes identified with different methods in human breast cancer research (the original study identified 14 marker genes). For **a)** and **b)**, intersections between gene identified from different methods were included in analyses. **c)** Gene enrichment analysis on SVGs identified by BSP on mouse cerebellum data using Slide-seq V2. **d)** Gene ontology enrichment analysis on SVGs identified in acute kidney injury studies. **e)** Distribution of cell type marker annotations from PanglaoDB on identified SVGs in mouse cerebellum study using Slide-seq V2. **f)** *Calb1* gene expression in the mouse cerebellum data using Slide-seq V2. The expression values were log-transformed, and those greater than 1.0 were normalized to 1.0. **g)** Expression and **h)** ISH of *Calb1* gene in an adult mouse brain, http://mouse.brain-map.org/experiment/show/75457491.

### 2.3 BSP accurately identifies SVGs in 2D space in biological studies

We applied BSP to four previously published 2D spatial transcriptomic datasets, including mouse olfactory bulb^11^ and human breast cancer obtained by ST sequencing^11^, hippocampus by SeqFISH^21^, and mouse hypothalamus preoptic region by MERFISH^22^. We followed the metric evaluation protocols proposed by SPARK and compared Identified SVGs with the provided marker genes in their original research^15^. The results were compared with SpatialDE, SPARK, and Spark-X. All the methods were run with the default parameters.

The mouse olfactory bulb dataset contains 11,274 genes measured on 260 spots using SRT sequencing. BSP detected 8 of 10 marker genes from the original study^11^, while SpatialDE detected 3, SPARK detected 8, and SPARK-X detected 0. The comparison between different methods is shown in **Figure 3A** and **Supplementary Figure 4**. The two marker genes that BSP missed are *Nmb* and *Sv2b*, with *p*-values of 0.0589 and 0.1806, respectively. We reason these missed marker genes have expression variances confined to many isolated, relatively small regions, which could result in the same variances in both big and small patches (**Supplementary Figure 5**). The human Breast cancer dataset contains 5,262 genes measured on 250 spots by SRT sequencing. BSP detected 12 of 14 marker genes identified as SVGs from the original study, while SpatialDE detected 7, SPARK detected 10, and SPARK-X detected 8. The result comparison is shown in **Figure 3B** and **Supplementary Figure 6**. The two marker genes that BSP missed were *PEG10* and *PIP*, with *p*-values of 0.1728 and 0.4371, respectively. The other two FISH-based datasets include the hippocampus dataset, consisting of 249 genes on 131 cells obtained by SeqFISH, and the mouse hypothalamus preoptic region composed of 160 genes on 257 cells by MERFISH. BSP identified most of the marker genes reported in the original studies. Detailed results for mouse olfactory bulb, human breast cancer, hippocampus, and hypothalamus preoptic regions are provided in **Supplementary Tables 2-5**, respectively.

**Figure 4:**
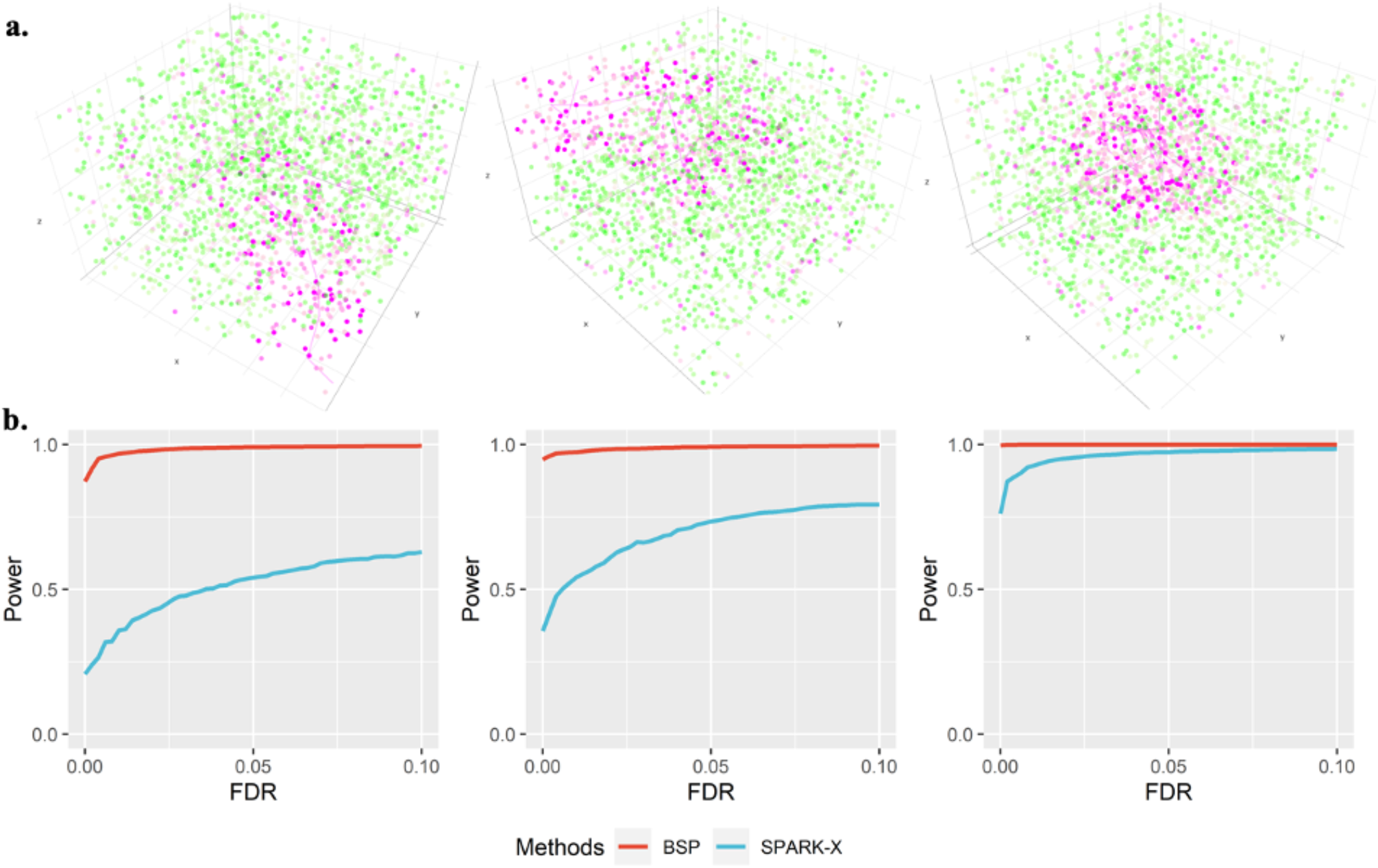
Power analysis for 3d simulation. **a)** Spatial patterns I-III are controlled by the direction of random walks. **Left**: Pattern I (curved stick): the movements of random walk are monotonic in two directions (x- and z-coordinates, or y- and z-coordinates). **Middle**: Pattern II (thin plate): the movements of the random walk are monotonic in one direction (z-coordinates). **Right**: Pattern III (irregular lump): the movements of the random walk are non-monotonic in any direction. **b)** Power comparison of the different methods under varied pattern sizes. Power charts show the averaged true positive rates (y-axis) across 10 replicates against the false discovery rates (x-axis) for the detected SVGs using each method. Simulations were performed using fixed moderate pattern sizes (*r* = 2.*0*), moderate signal strength (*FC* = 2.5), moderate noise level (*σ* = 1), and three spatial expression patterns I-III (left to right). All simulations were generated based on the seqFISH data with 10 segments (z-coordinate) and 225 spots of cells on each piece (x- and y-coordinates). Each simulation replicate contains 1,000 SVGs and 9,000 non-SVGs.

**Figure 5:**
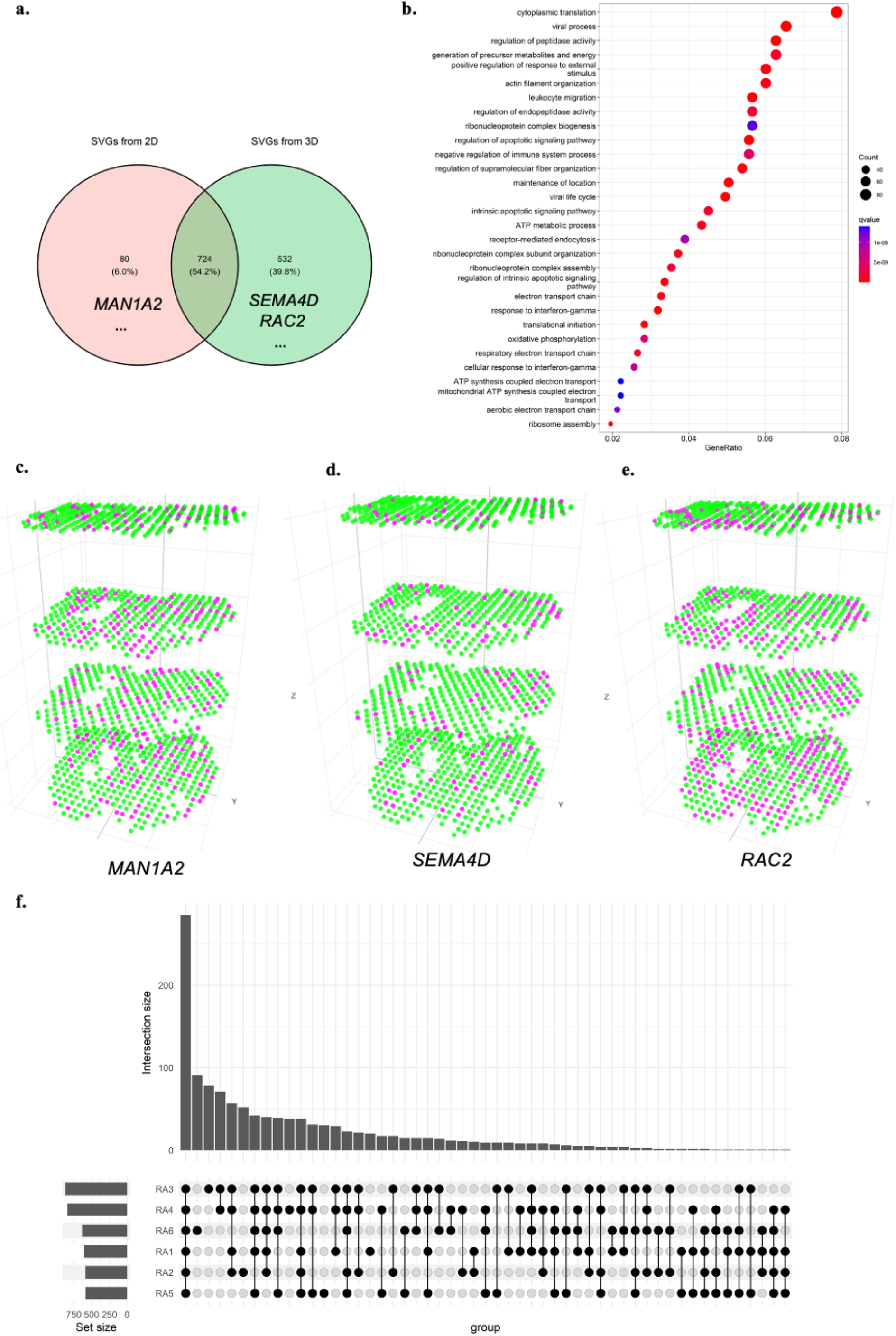
BSP identifies much more meaningful SVGs in the 3D study than stacking results on 2D analysis. **a)** Venn diagram of SVGs identified by 2D meta-analysis and 3D analysis; **b)** Gene ontology analysis on SVGs identified in patient RA1 in 3D setting; **c)** *MAN1A2* gene, significant in 2D analysis but not an SVG in 3D setting; **d)** *SEMA4D* gene identified as SVGs in 3D transcriptomics but missed by 2D analysis; **e)** *RAC2* gene, a very significant SVGs in 3D transcriptomics but missed by 2D analysis; **f)** Upset plot of enriched gene ontology terms on all six individual RA patients.

We extended the application of BSP in studies on Acute Kidney Injury (AKI)^23^. We ran BSP on five 10X Visium data on AKI samples collected in the Kidney Tissue Atlas^24^. BSP identified 285 SVGs (p-value<0.05) in one representative AKI sample consisting of 317 spots and 14,988 genes. Annotated by clusterProfiler^25^, the results were supported by a gene ontology (GO) enrichment analysis in **Figure 3C**, including relevant enrichments in humoral immune response (*q*-value 1.09E-11), and humoral immune response mediated by circulating immunoglobulin (*q*-value 1.63E-10). Reactome enrichment analysis^26^ identified eukaryotic translation elongation (*q*-value 2.77E-13) and influenza infection (*q*-value 5.59E-09), as in **supplementary Figure 7**. As innate and adaptive immune responses mediate damage to renal tubular cells and recovery from AKI^27,28^, these results are consistent with disease enrichment analysis^29^, which found the most significant terms in urinary system disease (*q*-value 9.30E-11) and kidney disease (*q*-value 1.36E-10). **Supplementary Table 6** lists all the SVG results obtained by BSP. **Supplementary Table 7** details the results from GO enrichment analysis, **Supplementary Table 8** details the results from Reactome, and **Supplementary Table 9** details the results from disease ontology enrichment analysis. By demonstrating BSP’s utility in kidney research, our study highlights the potential for BSP to advance our understanding of complex diseases in diverse tissue types.

### 2.4 BSP identifies SVGs on large-scale spatial transcriptomic studies using feasible computational resources

BSP was tested on three large-scale SRT datasets, including Slide-seq data on mouse cerebellum consisting of 18,671 genes on 25,551 beads, Slide-seq V2 data on mouse cerebellum consisting of 23,096 genes on 39,496 beads, and HDST olfactory bulb data consisting of 19,950 genes measured on 181,367 spots. BSP was successful on these large-scale datasets with a reasonable computational time. On the Ubuntu workstation described in Section 2.2, BSP took 7 and 18 minutes to process the Slide-seq mouse cerebellum and Slide-seq V2 mouse cerebellum data, respectively. The memory costs were around 19GB and 32GB, respectively. For the HDST olfactory bulb data, BSP took 4 hours and 90GB of memory on a High-Performance Computer equipped with Intel Xeon(R) CPU E5-2699 v4 @ 2.20GHz. The running details are listed in **Supplementary Table 10**.

BSP detected SVGs with a *p*-value less than or equal to 0.05 (n = 842, 1156, and 909 in Slide-seq V1 mouse cerebellum data, Slide-seq V2 mouse cerebellum data, and HDST olfactory bulb data, respectively), and we queried PanglaoDB^30^ with these detected SVGs. For each of the three implicated datasets, BSP returned numerous neuron-specific and non-neuron-specific genes. Detailed results of detected SVGs from each dataset are listed in **Supplementary Tables 11**-**13**. Interestingly, in addition to the SVGs corresponding to known cell type composition, many identified genes (30%) were not identified as any cell type markers with PanglaoDB annotations based on the knowledge from previous studies (**Figure 3E**).

On 1,156 identified SVGs in Slide-seq V2 (**Supplementary Table 14**), GO enrichment analysis showed significant enrichments in synapse organization (*q*-value 8.60e-51) (**Figure 3D, Supplementary Table 15)**. The expression patterns of five representative genes, *Calb1, Malat1, Nsg1, Ttc3*, and *Meg3*, were missed by SPARK-X and annotated as ‘unknown’ due to the low human brain regional specificity by the Human Protein Atlas^31^. *Calb1* gene (BSP *p*-value 2.73e-14, SPARK-X *p*-value 0.13, **Figure 3F**) is a Ca^2+^ buffering protein found to increase during postnatal development and decrease with aging and neurodegenerative disorders^32^. *Malat1* (BSP *p*-value 2.73e-14, SPARK-X *p*-value 0.63, **Supplementary Figure 8**) is a highly conserved nuclear-retained lncRNA shown to play a role in regulating genes at both the transcriptional and post-transcriptional levels in a context-dependent manner^33^. *Malat1* is shown to be dispensable for normal development and viability in mice^34^. *Ttc3* gene (BSP *p*-value 2.73e-14, SPARK-X *p*-value 0.30, **Supplementary Figure 9**) is known to play a role in cognitive impairment through protein quality control, which is a common phenotype of Down’s syndrome and Alzheimer’s disease^35^. Another representative gene *Nsg1* (BSP *p*-value 2.73e-14, SPARK-X *p*-value 1.00, **Supplementary Figure 10**), is known to be implicated in regulating endosomal recycling and sorting of several important neuronal receptors^36^. In addition, the *Meg3* gene (BSP *p*-value 2.73e-14, SPARK-X *p*-value 1.00, **Supplementary Figure 11**) modulates AMPA receptor surface expression in primary cortical neurons, and it is in the intricate regulation of the PTEN/PI3K/AKT signaling cascade during synaptic plasticity in neurons^37^. Besides Slide-seq V2, the spatial patterns of *Calb1, Ttc3, Nsg1*, and *Meg3* were validated by expression (**Figure 3G)** and ISH (**Figure 3H**) from Allen Brain Atlas^38^. Overall, the structural and functional compartmentalization in the cerebellum revealed by cell type annotation analysis highlights the utility of BSP.

### 2.5 BSP accurately identifies SVGs in 3D simulations

We extended the simulation framework in Trendsceek and SPARK further to demonstrate the power of BSP on 3D transcriptomic data. We compared the detection accuracy of SVGs using BSP with that of SPARK-X. The spatial patterns were constructed by a set of center points generated from a random walk with a fixed step length, and any spots within a certain distance from any of the center points were included as the marked cells. We created three 3D patterns, namely, curved stick (Pattern I), thin plate (Pattern II), and irregular lump (Pattern III), controlled by different directions of random walks, as shown in **Figure 4A**.

We performed the power analysis based on the FDR, considering the differences in the distribution of calibrated *p*-values. Compared to SPARK-X, BSP demonstrated superior and stable power under a wide range of FDR cutoffs with fixed moderate pattern sizes (*r* = 2.*0*), moderate signal strength (*FC* = 2.5), and moderate noise level (*σ* = 1) for the spatial expression patterns I, II, and III (**Figure 4B**). We also varied the pattern sizes, signal strengths, and noise levels while holding the other two parameters constant and found that BSP consistently demonstrated greater power in every scenario tested. **Supplementary Figure 12** shows the power analysis in different pattern sizes using a fixed moderate signal strength (2.5-fold) and low noise level (*σ* = 0). Simulations with pattern sizes as small (*r* = 1.5), moderate (*r* = 2.*0*), and large (*r* = 2.5) are tested on all patterns I, II, and III. **Supplementary Figure 13** demonstrates the results on different signal strengths using a fixed moderate pattern size (radius of 3) and low noise level (*σ* = 0). Simulations with signal strengths as low (2-fold), moderate (2.5-fold), and large (3-fold) are tested on patterns I, II, and III. **Supplementary Figure 14** shows the results on various noise levels using a fixed moderate pattern size (radius of 3) and moderate signal strength (3-fold). Simulations with high (*σ* = 2), moderate (*σ* = 1), and low (*σ* = 0) noise levels are tested on patterns I, II, and III.

### 2.6 BSP identifies more meaningful SVGs in the 3D study than stacking results on the 2D analysis

We utilized BSP on two publicly available 3D transcriptomics datasets, mouse visual cortex through STARmap sequencing^4^ and human RA synovium using stacking SRT^6^. The STARmap dataset contains 28 known SVGs (23 cell-type markers and 5 activity-regulated genes) measured in 33,598 spots. For these low throughput SRT with few genes, BSP adopted the generated null gene approach proposed by SPARK, and identified all these 28 genes as SVGs.

A study on human RA synovium contains 3D spatial transcriptomic sequencing from six RA patients by stacking 2D slices. Each sample consisted of approximately 13,000 genes on three to seven 2D slices with approximately 1,200 spots in each slice. To evaluate the power of 3D transcriptomics, BSP was first applied to each 2D slice, and then to the stacked 3D volume. Using the first sample (patient RA1) as an example, 260 genes were identified as the SVGs by taking the intersection of SVGs from four independent analyses on each 2D slice. However, 1,257 genes were detected as the SVGs by analyzing the stacked 3D SRT. All 260 genes from the 2D analysis were included in the gene list detected in 3D space, while 997 additional genes were discovered only in 3D space. We further examined these 997 genes neglected by 2D analysis with the DAVID functional annotations^39^ and found significant enrichments in host-virus interaction (Benjamini: *p*-value 4.3e-23), respiratory chain (Benjamini: *p*-value 3.4e-8), innate immunity (Benjamini: *p*-value 2.5e-6), neutrophil degranulation (Benjamini: *p*-value 7.1e-31), and viral process (Benjamini: 7.7e-17) among biological processes.

We also performed a classical meta-analysis by combining four individual analyses on each 2D slice using Fisher’s combined probability test with SciPy packages. The meta-analysis identified 804 genes as statistically significant (*p*-value < 0.05). Compared to the 1,257 SVGs identified by the 3D analysis, 724 genes were detected as SVGs by both the 2D meta-analysis and 3D settings (**Supplementary Table 16**), 532 genes were only significant in 3D settings (**Supplementary Table 17**), and 80 genes were only significant in the 2D meta-analysis setting (**Supplementary Table 18**). **Figure 5A** shows the Venn diagram of differences between meta-analysis and 3D analysis. **Figure 5B** shows GO enrichment analysis on all SVGs identified in 3D settings (**Supplementary Table 19**). Several immune-related gene ontologies are highlighted in RA studies, including response to interferon-gamma (*q*-value 2.25e-11)^40^, myeloid leukocyte migration (*q*-value 3.39e-09), leukocyte migration (*q*-value 3.45e-12), leukocyte chemotaxis (*q*-value 2.29e-09), regulation of leukocyte migration (*q*-value 5.11e-09)^41^. **Supplementary Figure 15** and **Supplementary Table 20** show GO enrichment results on 724 genes, both identified by 2D meta-analysis and 3D settings. The same GO enrichment analysis is proceeded on 532 genes uniquely identified by 3D settings in **Supplementary Figure 16** and **Supplementary Table 21**. These highlighted GO terms are indicative of key immune responses in immunizations in RA progression^42^.

2D meta-analysis may lead to some misleading results. Among these genes, *MAN1A2* (**Figure 5C**) gets Fisher’s combined *p*-value 7.56e-08 with four individual 2D *p*-values 0.8755, 6.83e-06, 0.0129, and 3.6e-04. However, the *p*-value of *MAN1A2* in 3D settings is 0.3648, making it unlikely as an SVG in 3D space when considering all the slices as a volume. On the other hand, among SVGs only significant in 3D analysis, *SEMA4D* plays a role in the immune system and induces B-cells to be aggregated and improves their viability (*in vitro*)^43^. Although the individual 2D *p*-values of *SEMA4D* are 0.9505, 0.9495, 0.9616, and 0.9735, its Fisher’s combined *p*-value in meta-analysis is 1.0, which has the least possibility of being an SVG in all individuals and meta-analysis on biased 2D analysis. However, the BSP test results of gene *SEMA4D* is 0.0482 on the 3D volume, making it stand out from the genes (**Figure 5D**). Another example that fails in the 2D analysis is *RAC2* (**Figure 5E**), which encodes a member of the Ras superfamily of small guanosine triphosphate (GTP)-metabolizing proteins involved in generating reactive oxygen species. Although Fisher’s combined *p*-value is 0.0942 with four individuals as 0.1173, 0.0846, 0.3286, and 0.3500, it is very significant with a *p*-value of 4.4929e-15 in the 3D setting. This analysis demonstrates that the analysis by BSP on intact 3D volume provides new opportunities in identifying SVGs compared to 2D analysis with potential bias.

The same analyses were conducted on each of the six RA patients individually. The enriched GO terms of 3D SVGs identified in each patient were presented in **Figure 5B** and **Supplementary Figures 17-21**. We observed that most GO terms were consistently enriched in all the patients (**Figure 5F**), indicating that BSP robustly identified 3D SVGs across various samples.

## 3. Discussion

The advancing spatial transcriptomics measures high-throughput multi-cellular- or cellular-level gene expression in the spatial context. This fast-growing 3D technology is critical for understanding the relationship between tissue structure and underlying biological function, posing new challenges in identifying SVGs vital in linking individual genes to spatial expression variance. The proposed BSP provides a dimension-agnostic and utilizes a big-small patch algorithm to identify SVGs at varying levels of granularity. The performance of BSP has been validated in both simulations and real studies using 2D and 3D data. While there is still a debate over the gold standard for SVGs in biological studies, we follow the protocol adopted by SPARK for analysis and annotation. Simulations provide an alternative benchmark for methods development. In the 2D simulation, BSP outperformed existing methods in most scenarios with different signal-to-noise ratios. In the 3D simulations, BSP demonstrated its superiority compared to other well-known criteria, such as Moran’s-I. Meanwhile, these 3D simulations can serve as benchmarks for developing new methods. In biological studies using 2D and 3D data, BSP identified more convincing SVGs than existing methods with a good control of false positives. For instance, in a human RA study, BSP revealed that analyzing SVGs as a volume in 3D data outperformed stacking results on individual 2D slices.

The innovation of BSP lies in its dimension-agnostic and granularity-guided approach, which utilizes paired big-small patches. Intuitively, the big patch provides a global view of the spatial pattern with a lower resolution, while the small patch focuses on the local details with a higher resolution. Using ratios between variances of the paired patches, BSP can accurately delineate the spatial patterns in a quantitative manner. Moreover, this approach is applicable to any dimension. These defined patches can effectively capture the characteristics of the expression patterns in both 2D space and 3D volume, making BSP capable of analyzing SRT data in both dimensions.

This granularity-guided approach makes the BSP a data-driven, model-free, and parameter-free model. First, BSP is particularly well-suited for the complexities of biological data, especially in the tumor microenvironment, where fixed spatial patterns cannot be assumed to form locally and globally. BSP’s effectiveness in these complex scenarios has been demonstrated in both 2D and 3D simulations, without preconceived assumptions about the underlying distributions. Second, BSP is robust to different levels of signal strengths and tolerates occasional noises, and it robustly discover the same persistent results in different samples, as the spatial pattens are invariant in different scale. Third, the BSP algorithm is highly efficient. In the typical scenario of a 10X Visium scale, BSP remains the fastest method among all the existing methods. Even for large-scale datasets, such as Slide-seq, Slide-seqV2, and HDST, BSP remains feasible with reasonable computational resources. Fourth, BSP’s core implementation is just a few dozen lines of code, making it easy to implement and adaptable to different usage scenarios.

Although BSP has shown significant improvements in quantitatively measuring spatial patterns using the beta distribution to fit the distribution of test scores of all the genes, some limitations still need to be addressed. Alternative statistical distributions or non-statistical ranking measurements could be explored to further improve the fitting of the distribution of ratios between variances of the averaged expression in the paired big-small patch. Furthermore, BSP compromises the performance and computational resources in SRT studies. Although gets superior performances on the benchmarks, BSP consumes more time and more memory than SPARK-X on large-scale datasets.

In conclusion, BSP has demonstrated its efficacy as a robust method for identifying SVGs in both 2D and 3D spatial transcriptomics analysis. As 3D sequencing technologies continue to advance and mature, we anticipate BSP to be increasingly valuable in future applications of 3D spatial transcriptomics. Moreover, as time is often considered as the fourth dimension in development biology^44,45^, we will also explore the potential for spatiotemporal studies using BSP.

## 4. Methods

### 4.1 BSP algorithm

BSP aims to identify spatially variable genes in 2D or 3D SRT data. The algorithm contains several steps, including (1) normalizing expression and spatial coordinates, (2) defining big and small patches for each spot based on neighboring spots with a larger or small radius, (3) calculating local means of gene expression for both big and small patches, (4) computing the ratio between the variances of local means between for big and small patches for each gene, and (5) fitting the ratio of each gene with a beta distribution and adjusting the p-values for each gene. The flowchart of the BSP algorithm is shown in **Supplementary Figure 22**.

#### 4.1.1 Problem setting and data normalization

On an SRT sample with *M* spots and *N* genes. The coordinates of spot *i* are (*x*_*i*_, *y*_*i*_) for 2D spatial transcriptomics, (*x*_*i*_, *y*_*i*_, *z*_*i*_) for 3D spatial transcriptomics. The expression level of gene *j* in spot *i* is denoted as 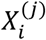, where 1 ≤ *i* ≤ *M*, 1 ≤ *j* ≤ *N*. The goal of BSP is to identify SVGs from all *N* genes with significant spatial patterns.All gene expression levels are normalized and scaled to [0,1] using a min-max normalization across all spots, 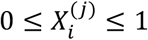 for all 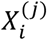. The normalization of spatial coordinates of the spots on SRT is based on the density of spots. The coordinates of spots in each direction are divided by the estimated density, which is calculated as the total number of spots divided by the area (2D) or volume (3D) of the sample. For simplicity, a rectangle is defined as the 2D space, and a cube is defined as the 3D volume. The rescaling functions *f* for 2D space is defined as Eq. (1), and for 3D space as Eq. (2):

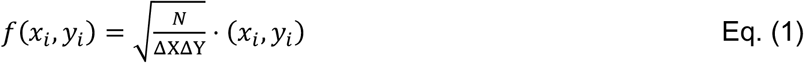

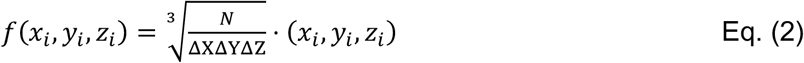

where *x*_*i*_, *y*_*i*_, and *z*_*i*_ are the coordinates of spot *i*. Δ*X*, Δ*Y*, and Δ*Z* denote the ranges of the sample space. They can be calculated as the differences between the maximum and minimum coordinates in each direction for the cube, as: Δ*X* = *max*(*x*) − *min*(*x*), Δ*Y* = *max*(*y*) − *min*(*y*), and Δ*Z* = *max*(*z*) − *min*(*z*).

This spatial coordinate normalization step ensures an adequate number of spots captured by the pre-defined radii *D*_*1*_ and *D*_*2*_. The goal of this step is to minimize the average spot-to-spot distance to slightly less than one unit. Typically, the default value of *D*_*1*_ is set as one unit to capture the nearest neighbors, while *D*_*2*_ is set to three units to include more spots in the patches.

#### 4.1.2 Big-small path

After coordinates normalization, the Euclidean distance between spots *i*_*1*_ and *i*_*2*_ is calculated as *dist*(*i*_*1*_, *i*_*2*_). For a given spot *i*, a **patch *S***_***i***_ is defined as the set of neighboring spots *l* within the radius of *D*

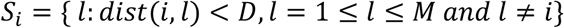

With a patch *S*_*i*_, 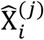 is defined as the **Local Mean**, the average expression level of gene *j* in this patch.

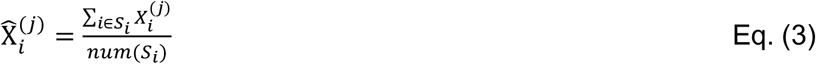

where *num*(*S*_*i*_) is the cardinal number of spots within *S*_*i*_. Local Mean describes the expression characteristics in the patch. 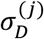 is defined as the variance of Local Means of all patches on gene *j*.

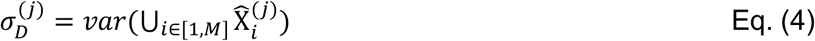

For a gene *j* without any spatial expression pattern, i.e., the distributions of the expression levels being identical across all spots, 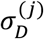 equals to 0. Otherwise, 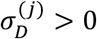 > 0. If distance *D* is big enough to cover the radius of the sample, then 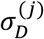 also equals 0 for each patch containing all spots, as the Local Means are the same for each spot.

For each spot *i*, we define a paired big-small patch, i.e., a mall patch is defined as *S*_*i*_′ with a radius *D*_*1*_, a big patch is defined as *S*_*i*_″ with a radius *D*_*2*_, where *D*_*1*_ < *D*_*2*_. We take the 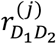, the ratio between the variances of the paired local averaged expression levels between big patch and small patch, describes the characteristics of the spatial pattern on gene *j*, defined as:

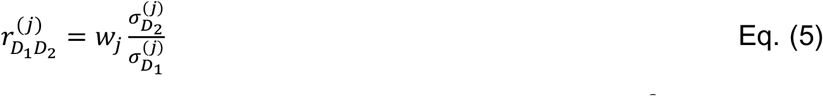

*w*_*j*_ is the weight to normalize the intrinsic gene expression variance, i.e., 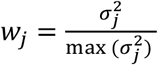, where 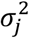 is the variance of expression levels of gene *j* of all the spots in the sample.

#### 4.1.3 Fitting beta distribution and adjusted p-value

After 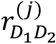 is calculated with all the genes, *j* ∈ [1, *M*], a beta distribution is approximated using the *stat* packages from sklearn^46^. If 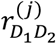 is greater than the 100 ∗ (1 − α)% upper tail of the fitted beta distribution, where *α* refers to the significance level (usually sets as 0.05), the significance p-value of each gene is addressed. In situations where there are insufficient null genes, a set of randomly permuted genes is generated to estimate the null distribution for practical usage.

### 4.2 2D simulation on mouse olfactory bulb data

We utilized the mouse olfactory bulb data within the framework of SPARK to construct 2D simulations. The simulation was based on mouse olfactory bulb data consisting of three spatial expression patterns measured on 260 spots (as shown in **Figure 2a**). Each simulation contained 1000 simulated SVGs with identified patterns in SpatialDE and SPARK, as well as 9000 non-SVGs generated through gene permutation without any spatial expression pattern. The p-values from basic spatial autocorrelation statistics Moran’s I, SpatialDE, SPARK, SPARK-X, and BSP were calculated to quantify the corresponding power (true positive rates) given a false discovery rate (FDR). To illustrate the rate of true positives (y-axis) identified by each method at different FDRs (x-axis) in power analysis, we generated ten replicates for each simulation. Specifically, simulation data was generated under different signal-noise ratios (*FC* = 3,4,5) with a medium level of noise (τ = 0.5, as defined in SPARK). Then another set of simulation data was generated under different noise levels (τ = 0.2,0.5,0.8) with a moderate signal-noise ratio (*FC* = 4).

### 4.3 3D simulation on FISH data

For the simulations in 3D space scenarios, we extended the framework originally introduced by Trendsceek and SPARK. All simulations were generated based on seqFISH data, with 10 segments in the z-coordinate and 225 spots representing cells in each piece in the x- and y-coordinates. We assume the sample was cryosectioned into 10 sections, with each section placed on an individual array without any direct contact between array surfaces. To generate spatial locations for a fixed number of cells (*n* = 225) in each section, we used a random-point-pattern Poisson process. These spatial locations for each section were then stacked together with the index of the section serving as the z-coordinates (*z* = 1,2, *…*, 10).

The 3D spatial patterns were constructed using a set of spheres with center points generated through a random walk with a fixed step length of 2. We included three types of spatial patterns in the simulations by controlling the range of directions (**Figure 4a**). These patterns include Pattern I (curved stick), the movements of a random walk are monotonic in two directions (x- and z-coordinates, or y- and z-coordinates); Pattern II (thin plate), the movements of a random walk are monotonic in one direction (z-coordinates); Pattern III (irregular lump), the movements of a random walk are non-monotonic in any directions. We produced 1000 SVGs with 3D patterns for each simulation, and generated 9000 non-SVGs without any spatial expression pattern by permutating known patterns.

The expression of SVGs was sampled based on whether the cell was inside or outside the pattern, distinguishing between marked and non-marked cells. For marked cells inside the pattern, we randomly selected gene expression values from the upper quantile of the gene expression distribution in the seqFISH data. For non-marked cells and those outside the pattern, we assigned gene expression randomly from the expression measurements in the seqFISH data. Non-SVGs were generated by permutating gene expressions of SVGs. For each SVG, the expression values were permuted and repeated 9 times (i.e., randomly assigning values to all cells without replacement). Finally, random noise was added proportionally to the averaged standard deviation of expressions in all genes.

To systematically explore the influences under different scenarios, we held two parameters constant while manipulating the third to vary the patterns’ sizes, signal strengths, and noise levels. We tested three sphere radius values (*r*) of 1.5, 2.0, and 2.5, which determined the pattern size. Quantile thresholds of 0.66, 0.80, and 0.88 were set, corresponding to expected expression fold changes (*FC*) of 2, 2.5, and 3 between marked cells and non-marked cells, indicating low, moderate, and strong signal strengths, respectively. We applied random noise following a Gaussian distribution with mean zero and the standard deviation (*σ*) of 0, 1, and 2 times the averaged standard deviation of the expressions of all simulated genes to represent low, moderate, and high noise levels. In the 3D simulations we varied the pattern sizes (*r* = 1.5,2.*0*,2.5), expression fold changes (*FC* = 2,2.5,3), and noise levels (*σ* = 0,1,2) across spatial patterns I, II, and III. For each combination of pattern size, signal strength, and noise level, we conducted 10 replicates to perform the power analysis.

### 4.4 Biological data collection and analysis

For studies on mouse olfactory bulb, human breast cancer obtained by SRT sequencing, hippocampus by SeqFish, and mouse hypothalamus preoptic region by MERFISH, we followed the analysis protocol adopted by SPARK. For studies on Slide-seq data, Slide-seqV2 data, HDST data, we followed the analysis protocol adopted by SPARK-X.

In the case of human RA synovium studies, the spatial locations in 2D slices were normalized with unit one. These 2D slices were stacked together with interval one on the z-axis to construct a volume on 3D transcriptomics. Analysis was performed based on the normalized data provided by the authors.

For kidney analysis, all the data were generated using 10X Visium platforms and processed with CellRange. Expression data is quality controlled and preprocessed by Seurat with scTransform^47^.

### 4.5 Annotations

The annotations using PanglaoDB were performed by rPanglaoDB (https://github.com/dosorio/rPanglaoDB). Go enrichment analysis was performed by topGO^48^ (Version 3.16). Reactome pathway analysis was performed by ReactomePA^26^ (Version 3.16). Disease Ontology Semantic and Enrichment analysis was performed by DOSE^29^ (Version 3.16). Meta-analysis is performed by SciPy from the sklearn package^46^ (Version 1.1.2) in python 3.9.12.

## Data Availability

The mouse olfactory bulb and human breast cancer data are available at http://www.spatialtranscriptomicsresearch.org, the MERFISH data can be downloaded from (https://datadryad.org/stash/dataset/doi:10.5061/dryad.8t8s248), and the SeqFISH data is available at https://www.cell.com/cms/10.1016/j.neuron.2016.10.001/attachment/759be4dc-04a6-4a58-b6f6-9b52be2802db/mmc6.xlsx. Slide-seq data, Slide-seqV2 data, HDST data, and human rheumatoid arthritis synovium data are available at Broad Institute’s single-cell repository (https://singlecell.broadinstitute.org/single_cell/) with ID SCP354, SCP948, SCP420, and SCP1414. The STARmap data set is available at https://www.starmapresources.com/data. The kidney spatial transcriptomics data can be downloaded from the Kidney Tissue Atlas (https://atlas.kpmp.org/). The generated simulation data is available at https://mailmissouri-my.sharepoint.com/:f:/g/personal/wangjue_umsystem_edu/EnjH6hbt1ptBjWBzoQhGfPoBhFVVynkeJbrdpszmeqnYpA?e=jDHmEJ, and will be deposited in the Zenodo database.

## Code Availability

The source code of BSP is freely available at https://github.com/juexinwang/BSP/.

## Acknowledgments

This work was supported by grants R35-GM126985 and R01-GM131399 from the National Institutes of Health, as well as the Pelotonia Institute of Immuno-Oncology (PIIO).

