## Supplementary Figures and Tables for "Dimension-agnostic and granularity-based spatially variable gene identification": Supplement_Figures.pdf

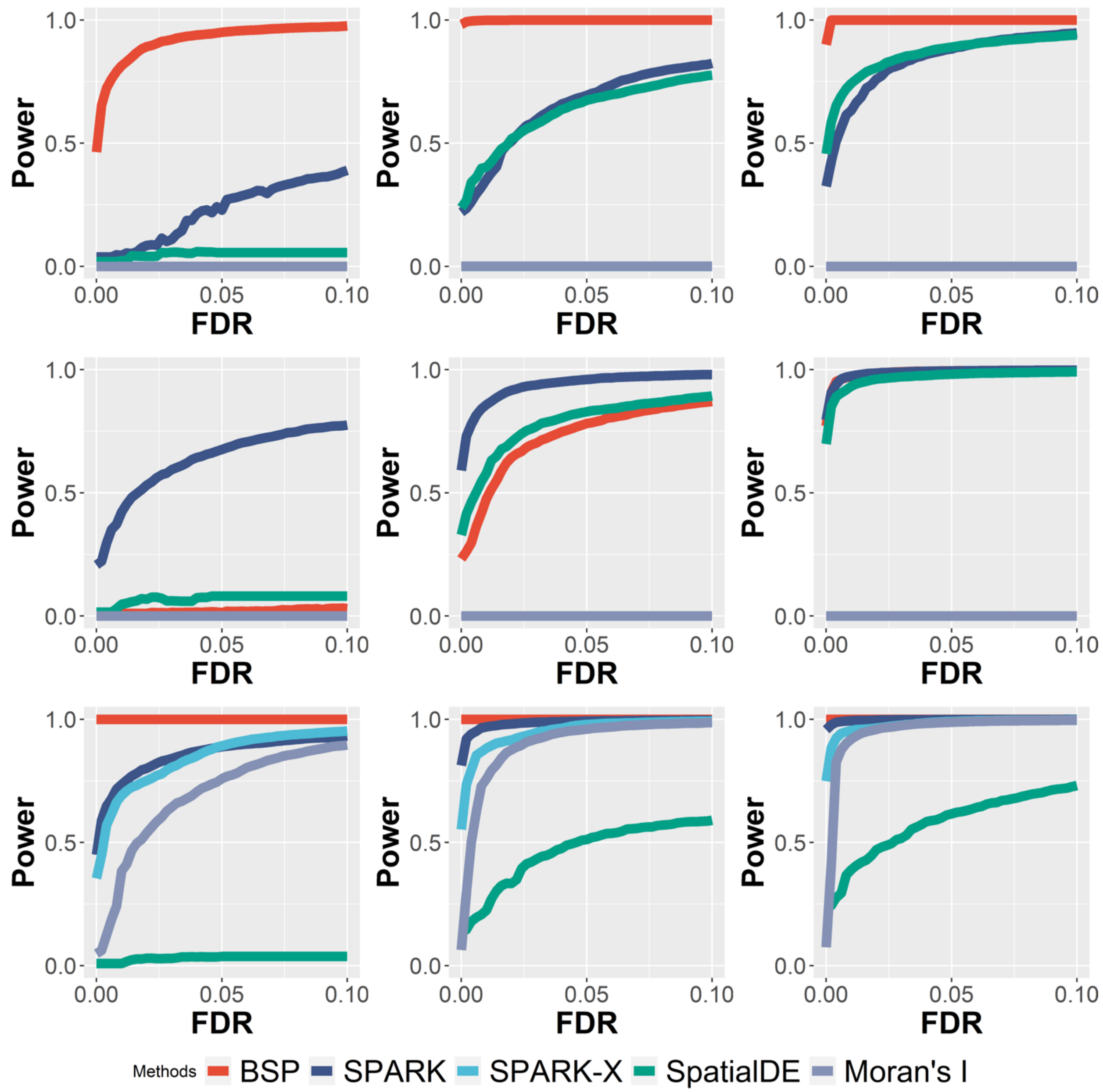


**Supplementary Figure 1. Power comparison of different methods under varying signal strengths in 2D simulation.** Power charts show the averaged true positive rates (y-axis) across 10 replicates against the false discovery rates (x-axis) for the detected SVGs using each method. In these nine power charts, simulations with low (2-folds), moderate (3-folds), and high (4-folds) signal strengths are shown in the left, middle and right columns, respectively. Simulations using the spatial expression patterns I, II, and III are placed in the top middle, and bottom rows, respectively. All simulated datasets were generated using a fixed noise level ($\tau$=0.5).

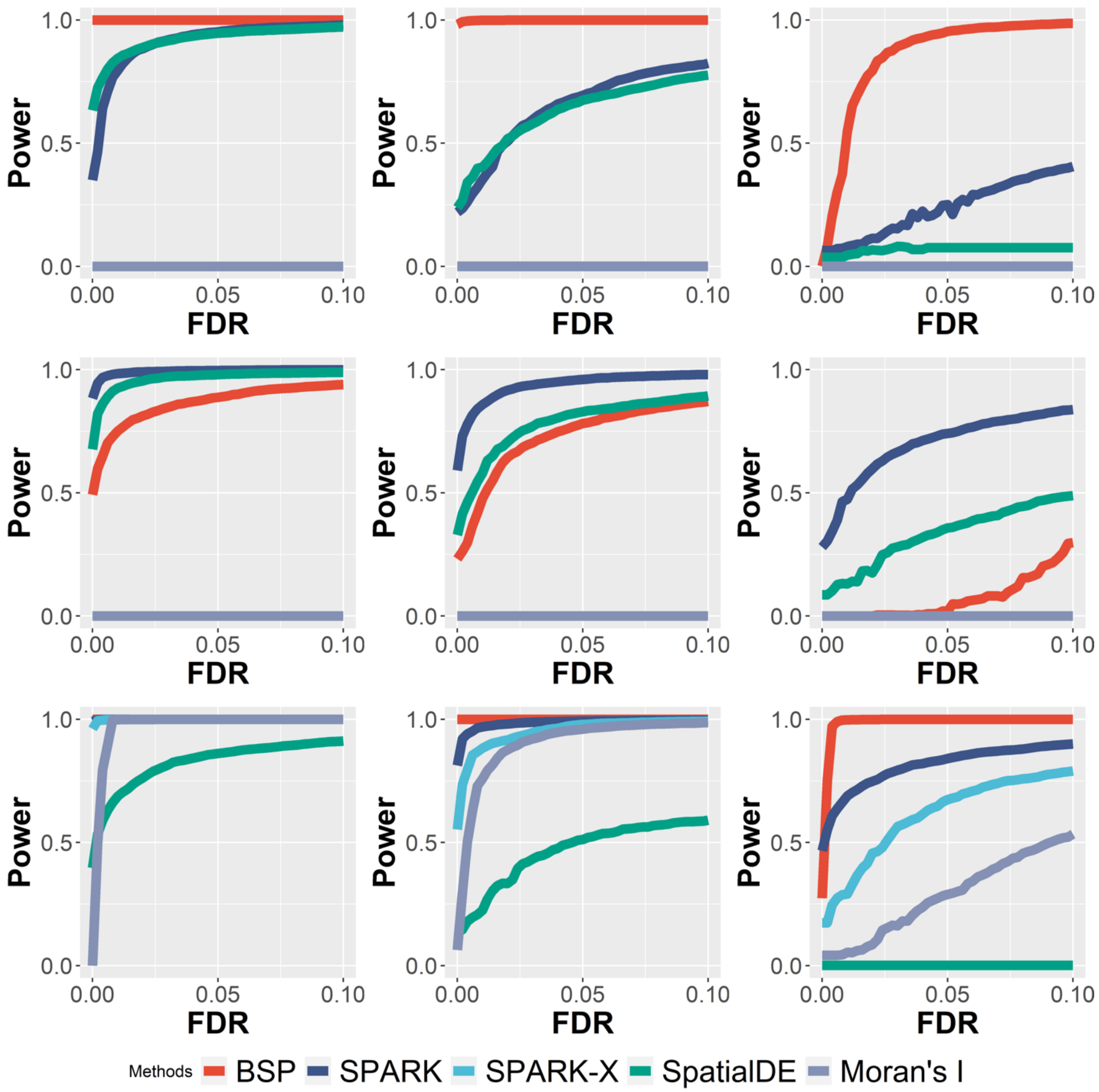


**Supplementary Figure 2: Power comparison of different methods under varying noise levels in 2D simulation.** Power charts show the averaged true positive rates (y-axis) across 10 replicates against the false discovery rates (x-axis) for the detected SVGs using each method. In these nine power charts, simulations with low ($\tau = 0.2$), moderate ($\tau = 0.5$), and high ($\tau = 0.8$) noise levels are shown in the left, middle and right columns, respectively. Simulations using the spatial expression patterns I, II, and III are placed in the top middle, and bottom rows, respectively. All simulated datasets were generated using a fixed moderate signal strength (3-folds).

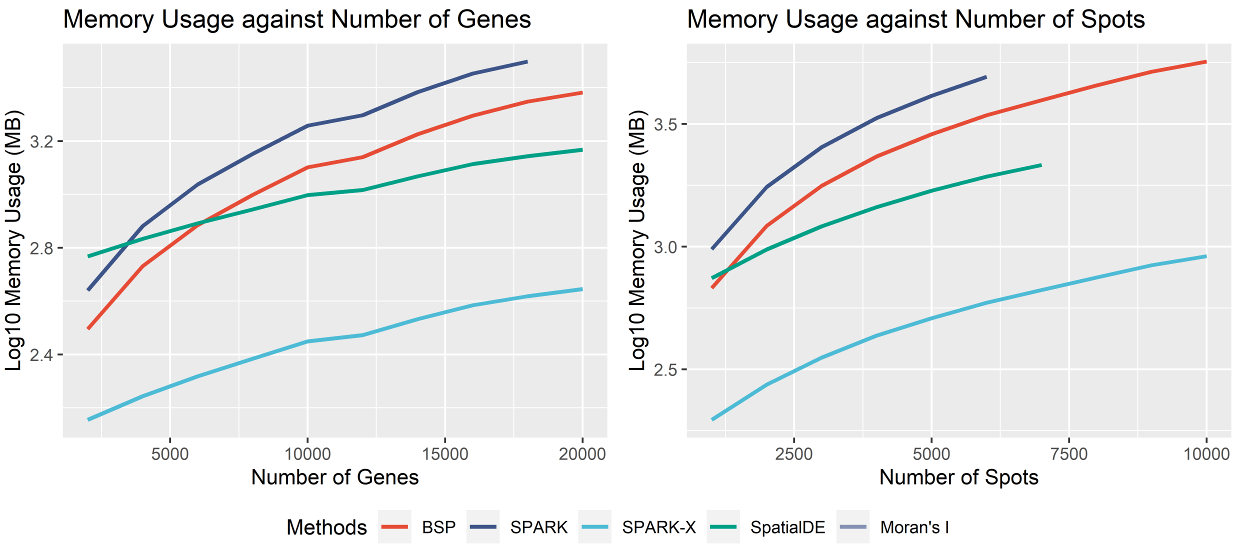


**Supplementary Figure 3. BSP memory usage with different gene numbers and different numbers of spots.** The experiment is performed on an Ubuntu 16.04.4 LTS workstation with Intel(R) Xeon(R) W-2125 CPU @ 4.00GHz and 32 GB memory.

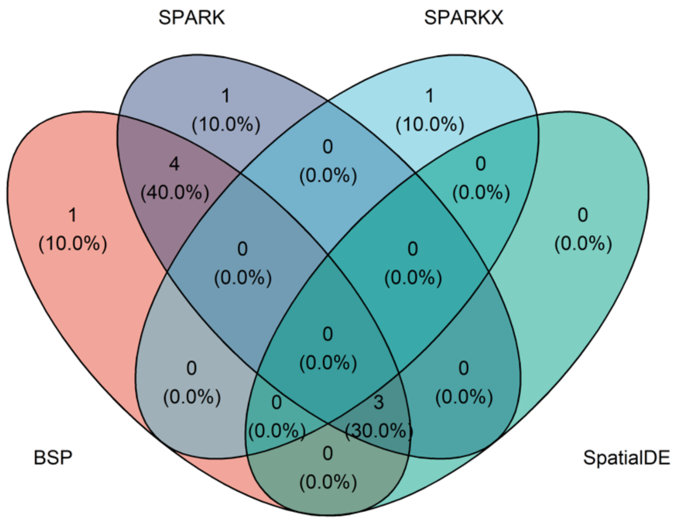


**Supplementary Figure 4. Venn diagram of marker genes identified with different methods in mouse olfactory bulb research**. The original study includes 10 marker genes.

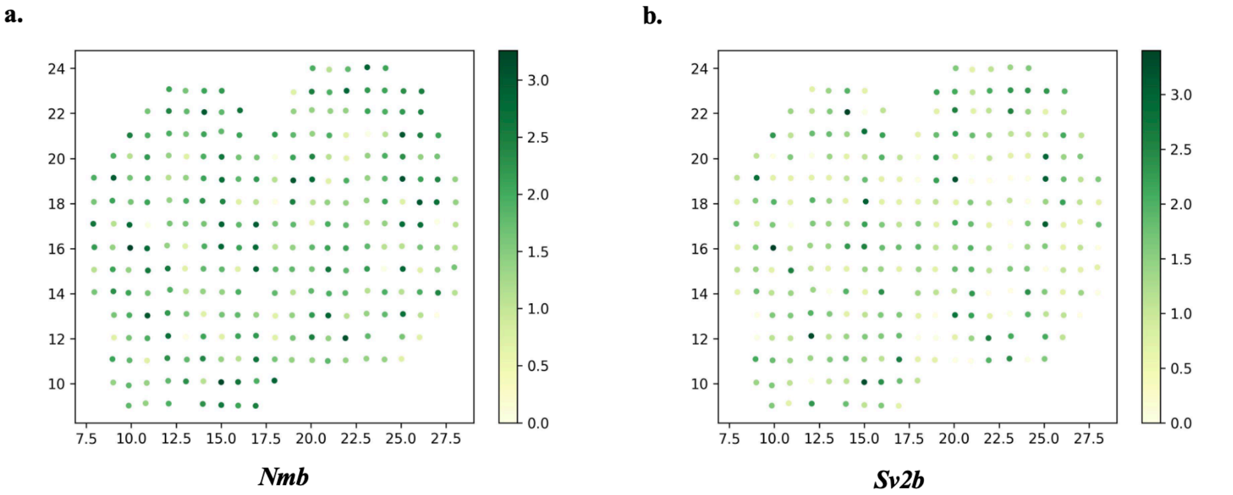


**Supplementary Figure 5. Missed marker genes by BSP in mouse olfactory bulb study**. a) Gene *Nmb*; b) Gene *Sv2b*. Colors indicate gene expression levels. The expression values were log-transformed, and those greater than 1.0 were normalized to 1.0.

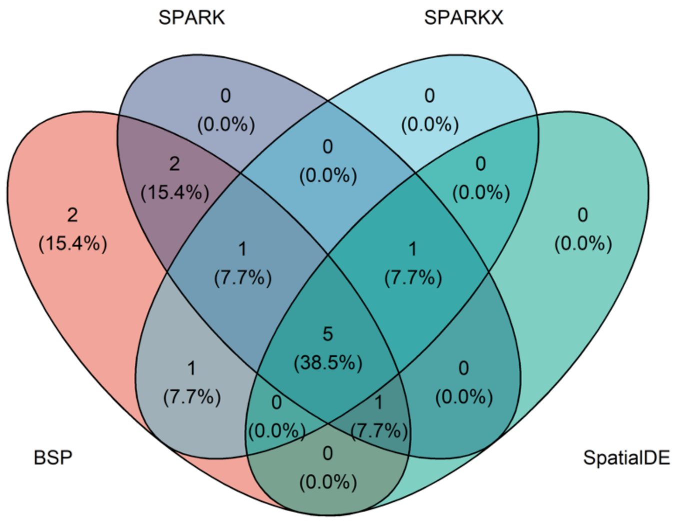


**Supplementary Figure 6. Venn diagram of marker genes identified with different methods in human breast cancer research**. The original study identified 14 marker genes.

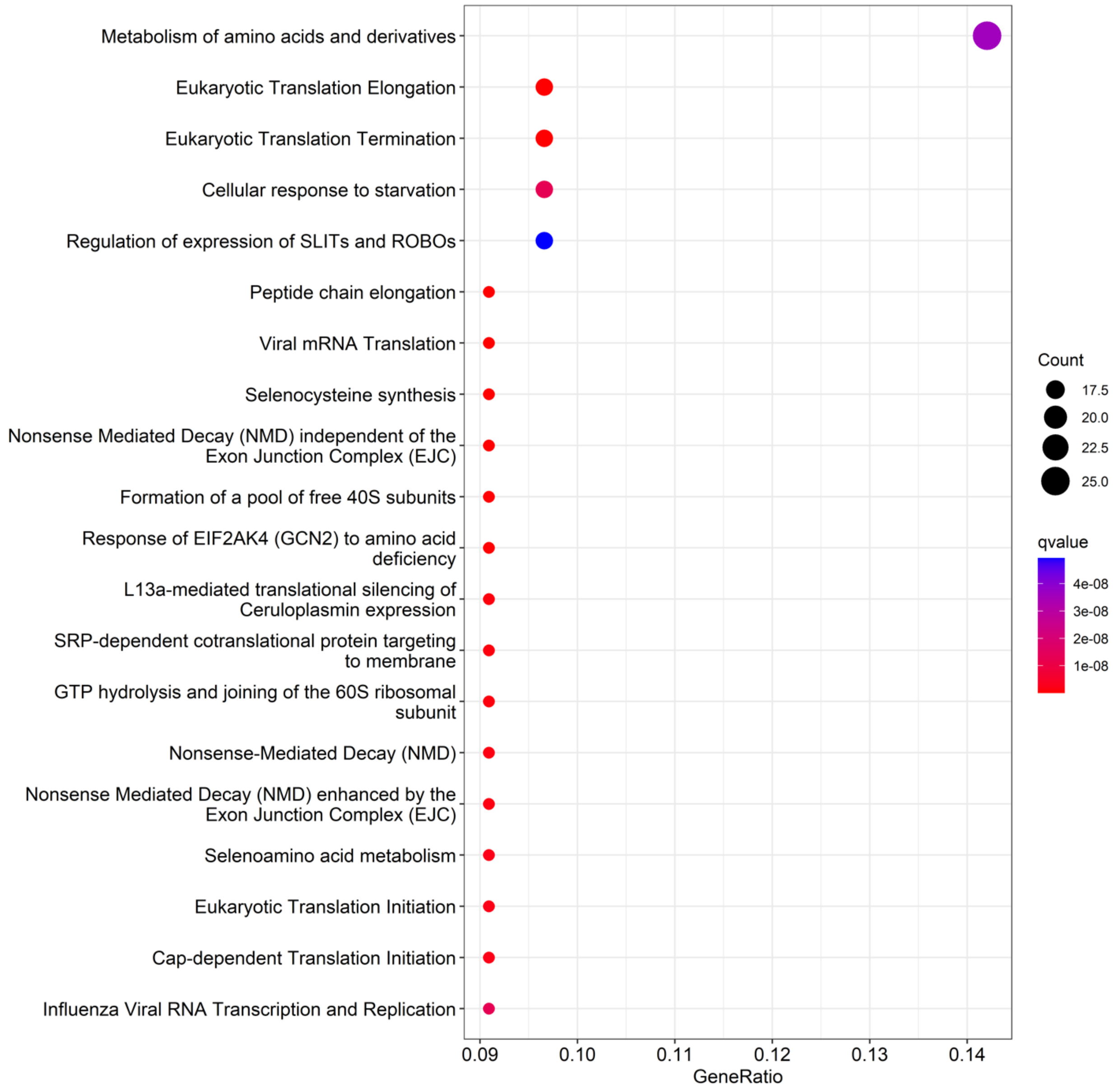


**Supplementary Figure 7: Pathway enrichment analysis on SVGs in AKI study using 10X Visium.**

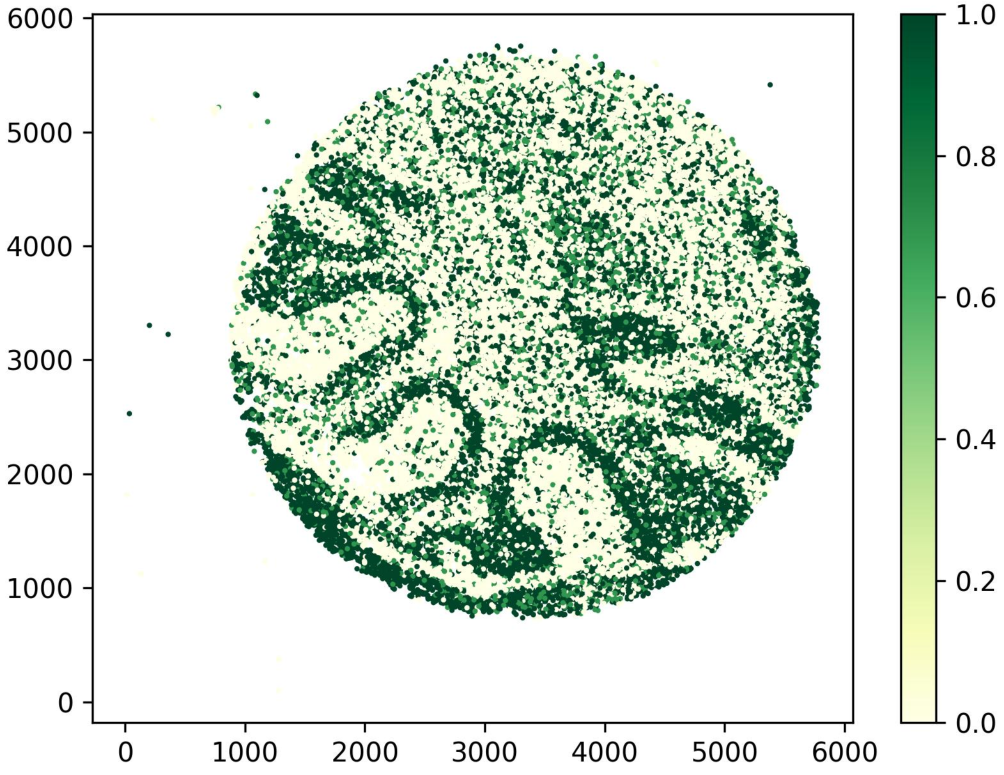


**Supplementary Figure 8. Showcase of *Malat1* as SVG identified by BSP.** Allen Brain Atlas does not have expression or ISH available at mouse.brain-map.org. Colors indicate gene expression levels. The expression values were log-transformed, and those greater than 1.0 were normalized to 1.0.

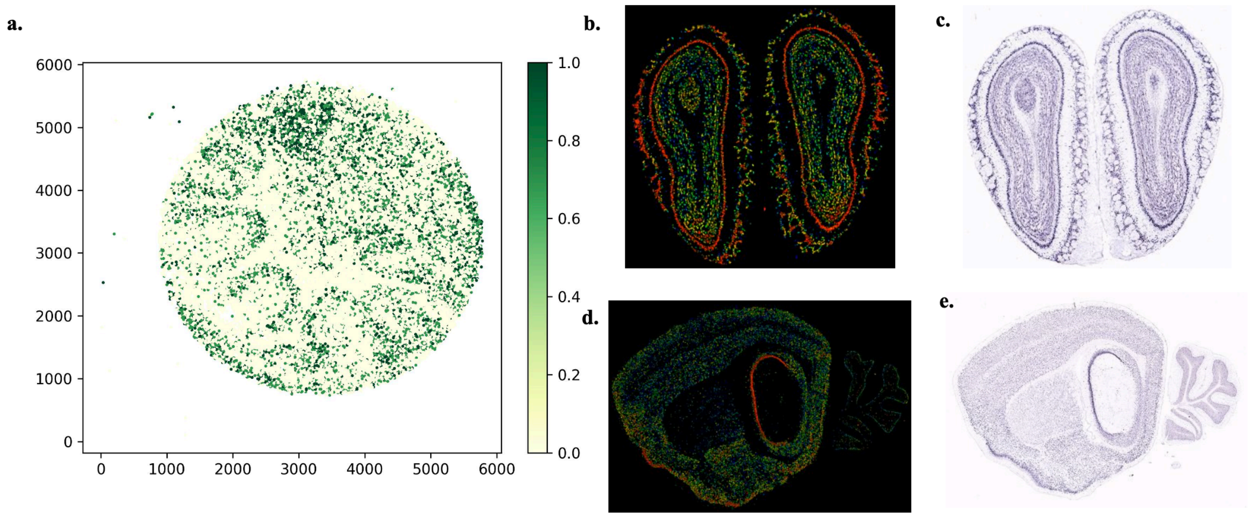


**Supplementary Figure 9. Showcase of *Ttc3* as SVG identified by BSP. a)** *Ttc3* gene expression in the mouse cerebellum data using Slide-seq V2; **b)** Expression and **c)** ISH of *Ttc3* gene in coronal adult mouse brain http://mouse.brain-map.org/experiment/show/1079; **d)** Expression and **e)** ISH of *Ttc3* gene in an sagittal adult mouse brain http://mouse.brain-map.org/experiment/show/1080

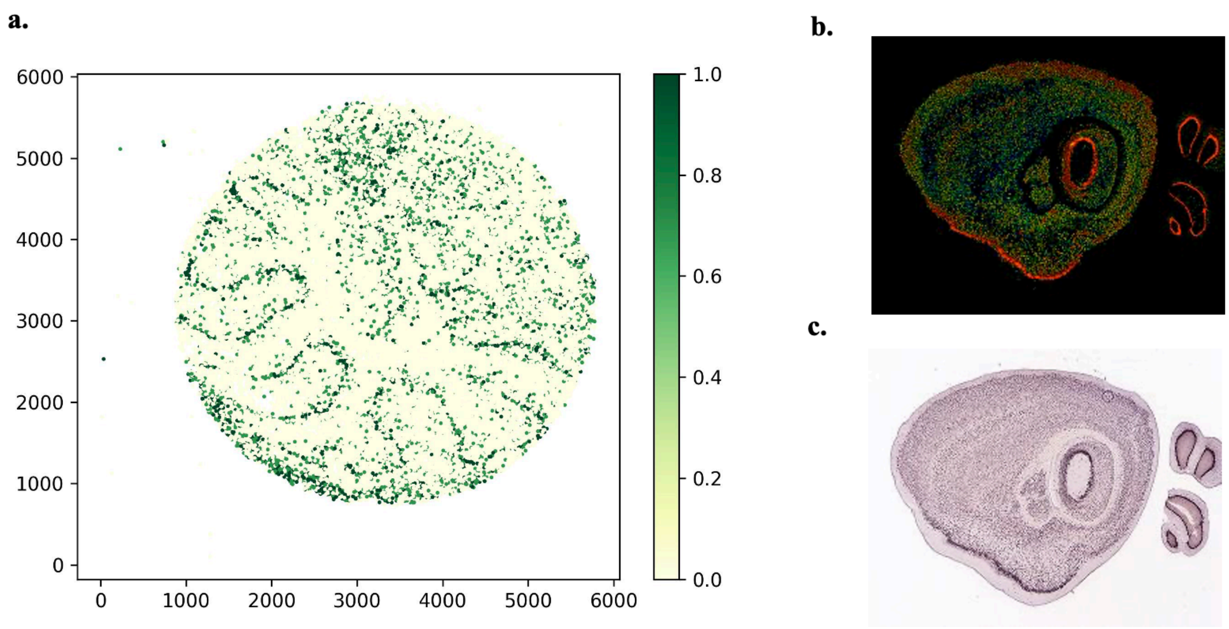


**Supplementary Figure 10. Showcase of *Nsg1* as SVG identified by BSP**. **a)** *Nsg1* gene expression in the mouse cerebellum data using Slide-seq V2; **b)** Expression and **c)** ISH of *Nsg1* gene in sagittal adult mouse brain [http://mouse.brain-map.org/experiment/show/70429327](http://mouse.brain-map.org/experiment/show/70429327)

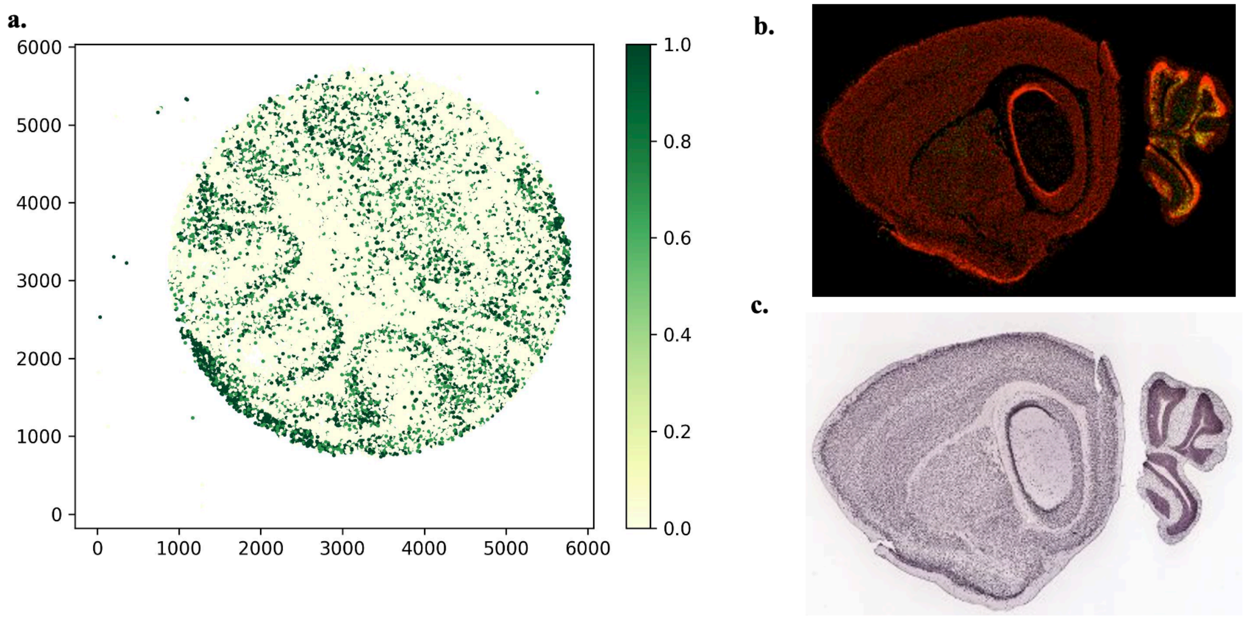


**Supplementary Figure 11. Showcase of *Meg3* as SVG identified by BSP. a)** *Meg3* gene expression in the mouse cerebellum data using Slide-seq V2; **b)** Expression and **c)** ISH of *Sparcl1* gene in sagittal adult mouse brain http://mouse.brain-map.org/experiment/show/71281027

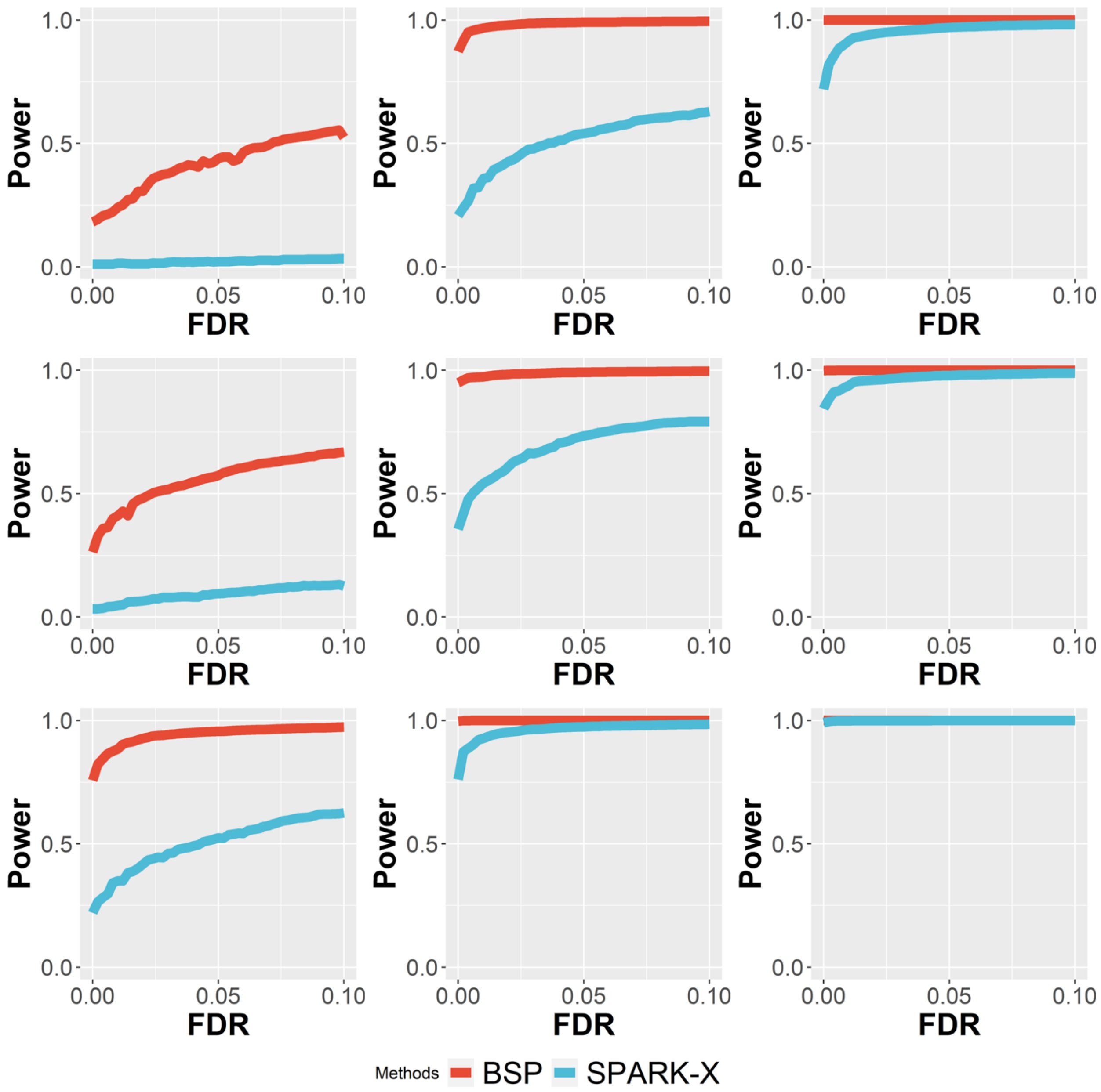


**Supplementary Figure 12: Power comparison under varying pattern sizes in 3D simulations**. Simulations were performed using a fixed moderate signal strength (2.5-fold) and low noise level ($\sigma = 0$). In these nine power charts, simulations with pattern sizes as small ($r = 1.5$), moderate ($r = 2.0$), and large ($r = 2.5$) are in the left, middle, and right columns, respectively. Simulations using the 3D Pattern I (curved stick), pattern II (thin plate), and pattern III (irregular lump) as in the top, middle, and bottom rows, respectively.

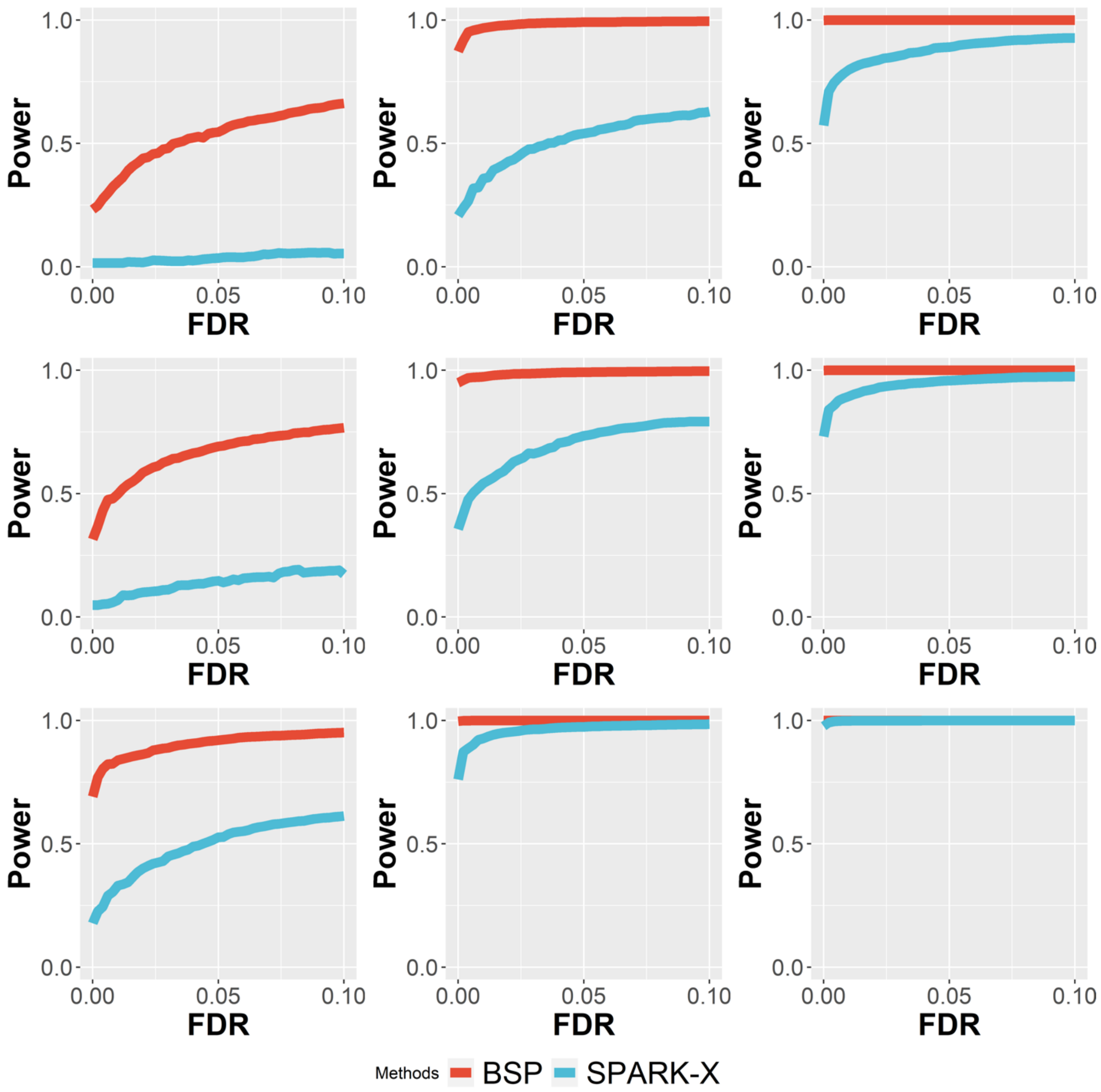


**Supplementary Figure 13: Power comparison under varying signal strengths in 3D simulations.** Simulations were performed using a fixed moderate pattern size (radius of 3) and low noise level ($\sigma = 0$). In these nine power charts, simulations with signal strengths as low (2-fold), moderate (2.5-fold), and large (3-fold) are in the left, middle and right columns, respectively. Simulations using the 3D Pattern I (curved stick), pattern II (thin plate), and pattern III (irregular lump) are in the top, middle, and bottom rows, respectively.

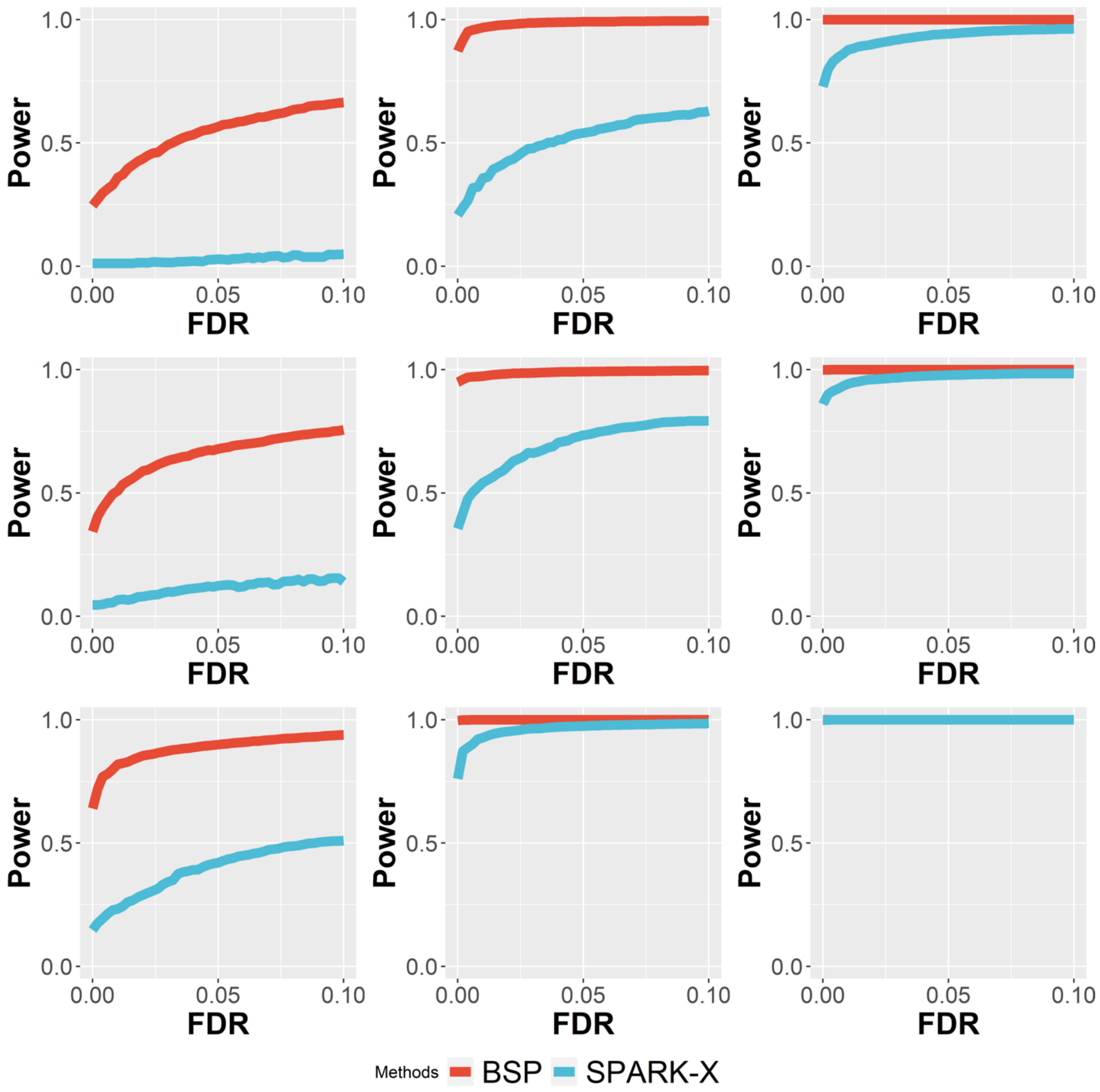


**Supplementary Figure 14: Power comparison under varying noise levels in 3D simulations.** Simulations were performed using a fixed moderate pattern size (radius of 3) and moderate signal strength (3-fold). In these nine power charts, simulations with high ($\sigma = 2$), moderate ($\sigma = 1$), and low ($\sigma = 0$) noise levels are in the left, middle, and right columns, respectively. Simulations using the 3D Pattern I (curved stick), pattern II (thin plate), and pattern III (irregular lump) are shown in the top, middle, and bottom rows, respectively.

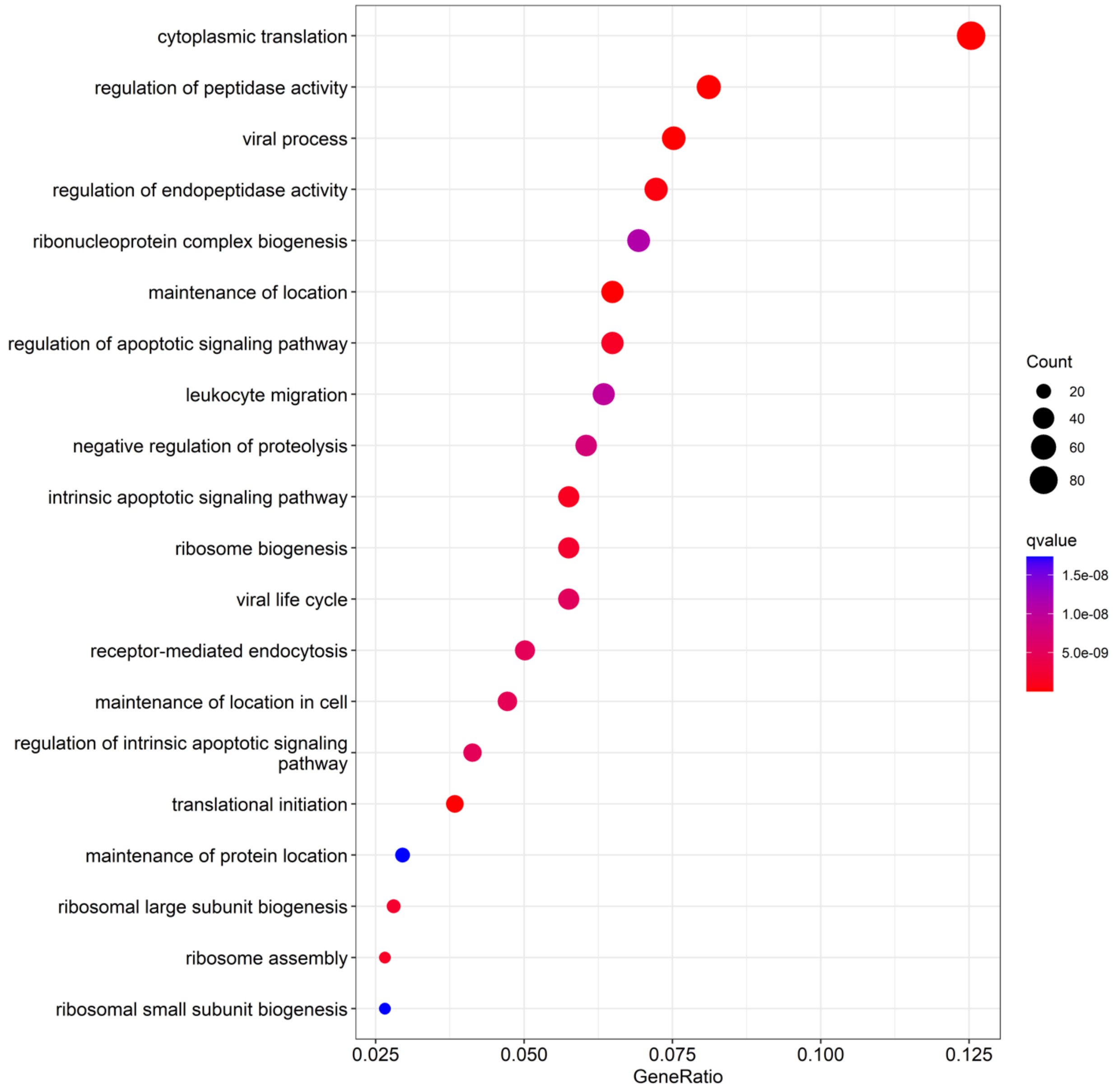


**Supplementary Figure 15: Gene ontology enrichment analysis on 724 genes both identified by 2D meta-analysis and 3D settings in patient RA1.**

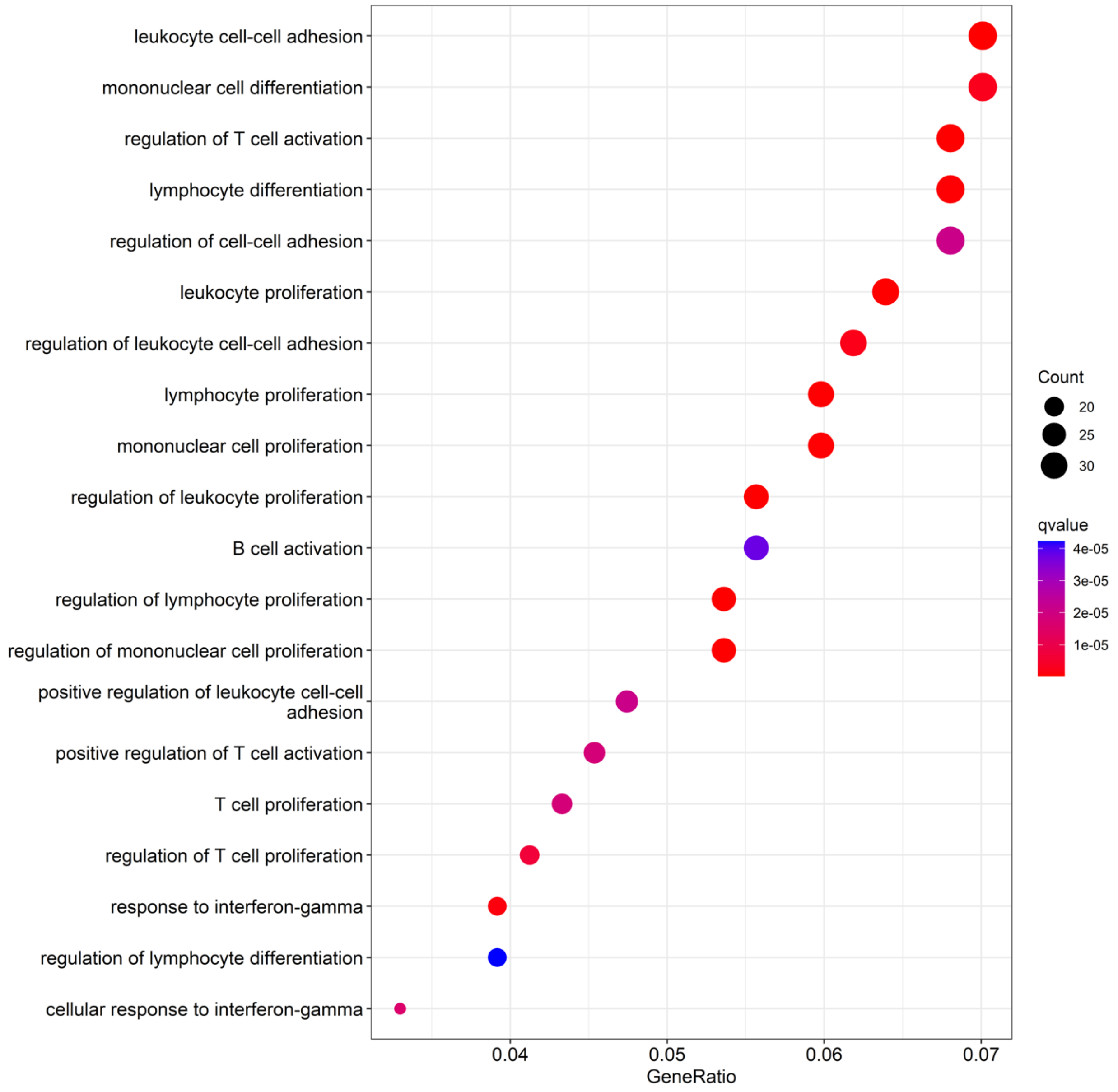


**Supplementary Figure 16: Gene ontology enrichment analysis on 532 genes uniquely identified by 3D settings in patient RA1.**

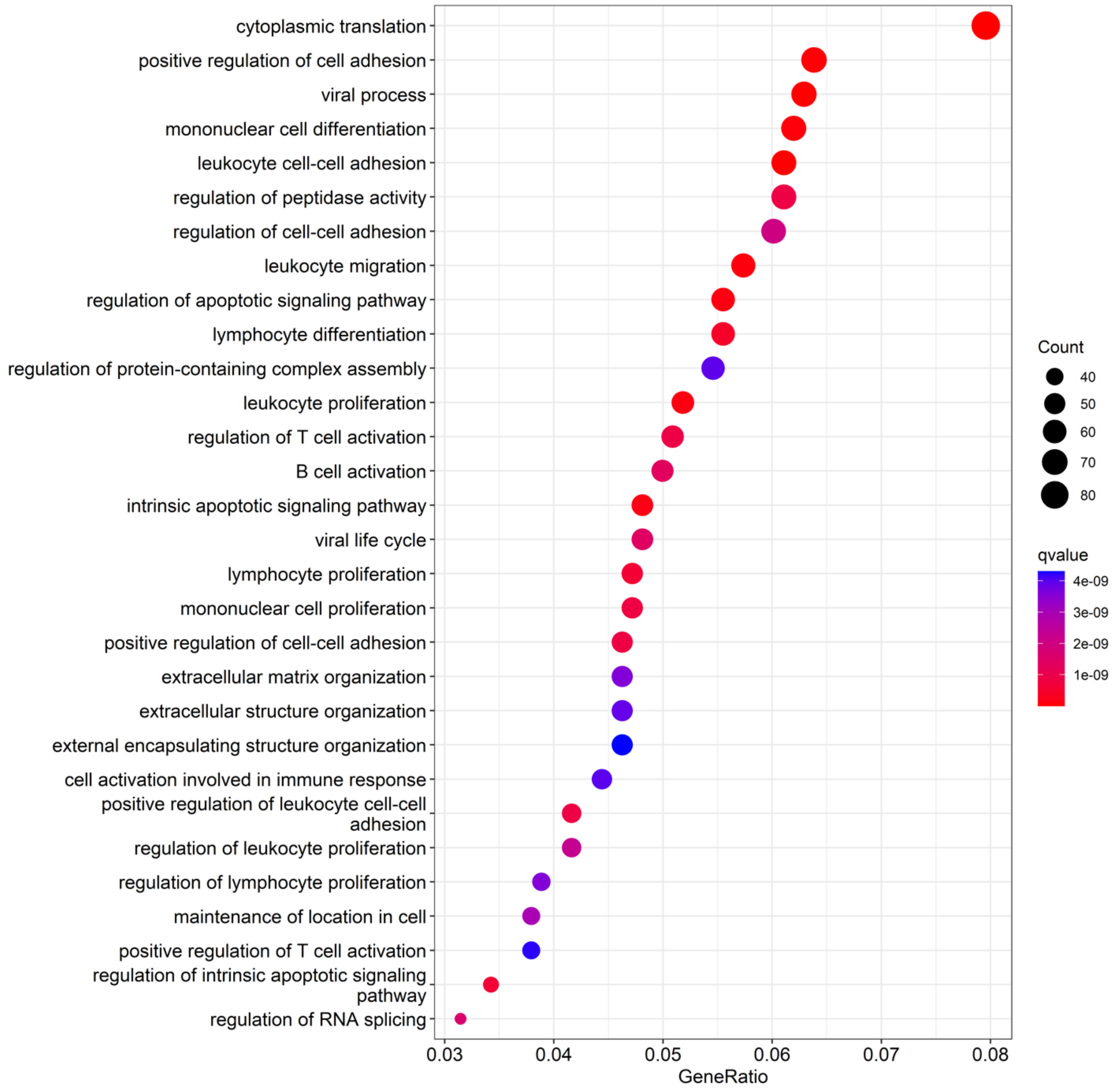


**Supplementary Figure 17: Gene ontology enrichment analysis on patient RA2.**

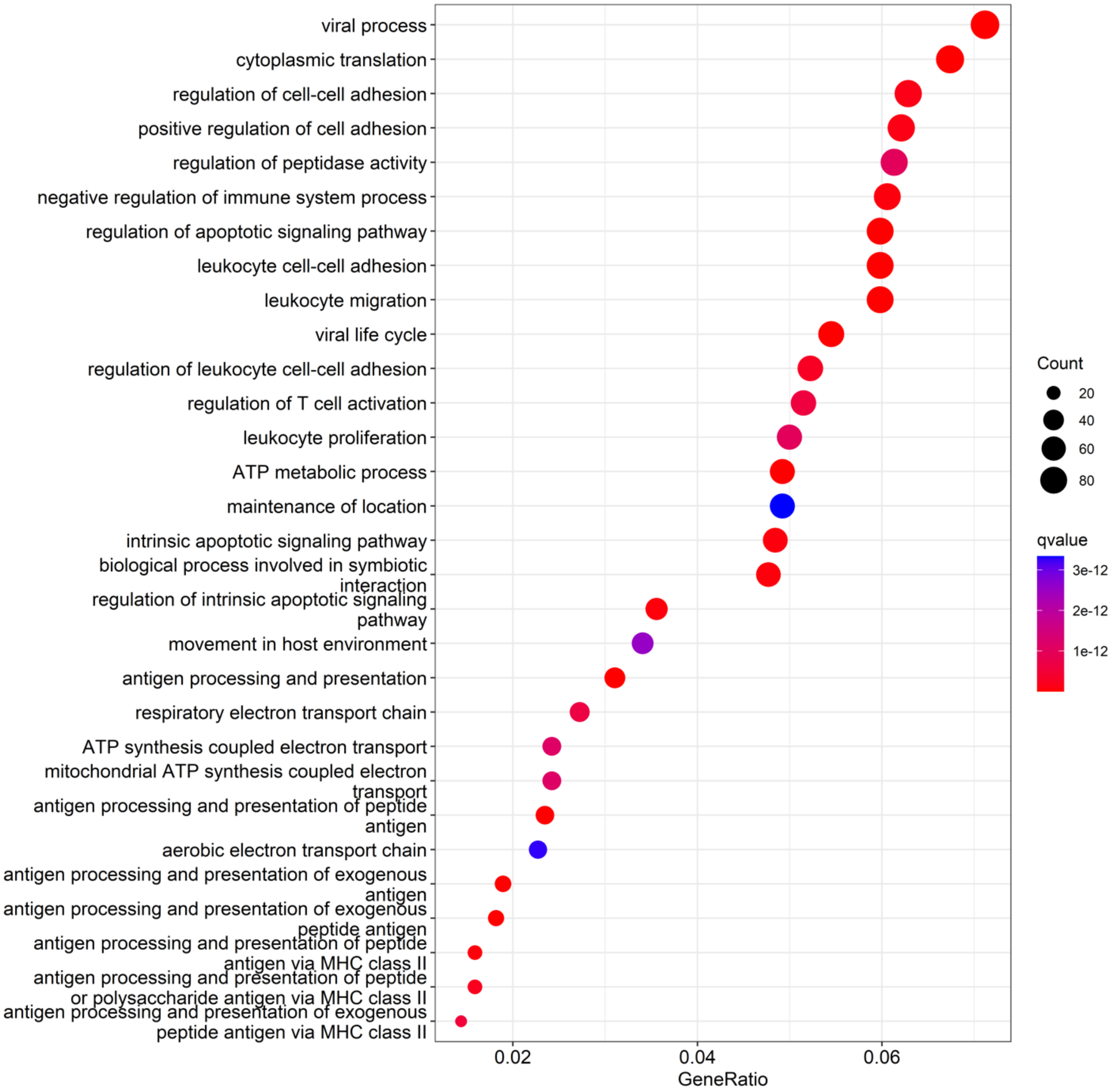


**Supplementary Figure 18: Gene ontology enrichment analysis on patient RA3.**

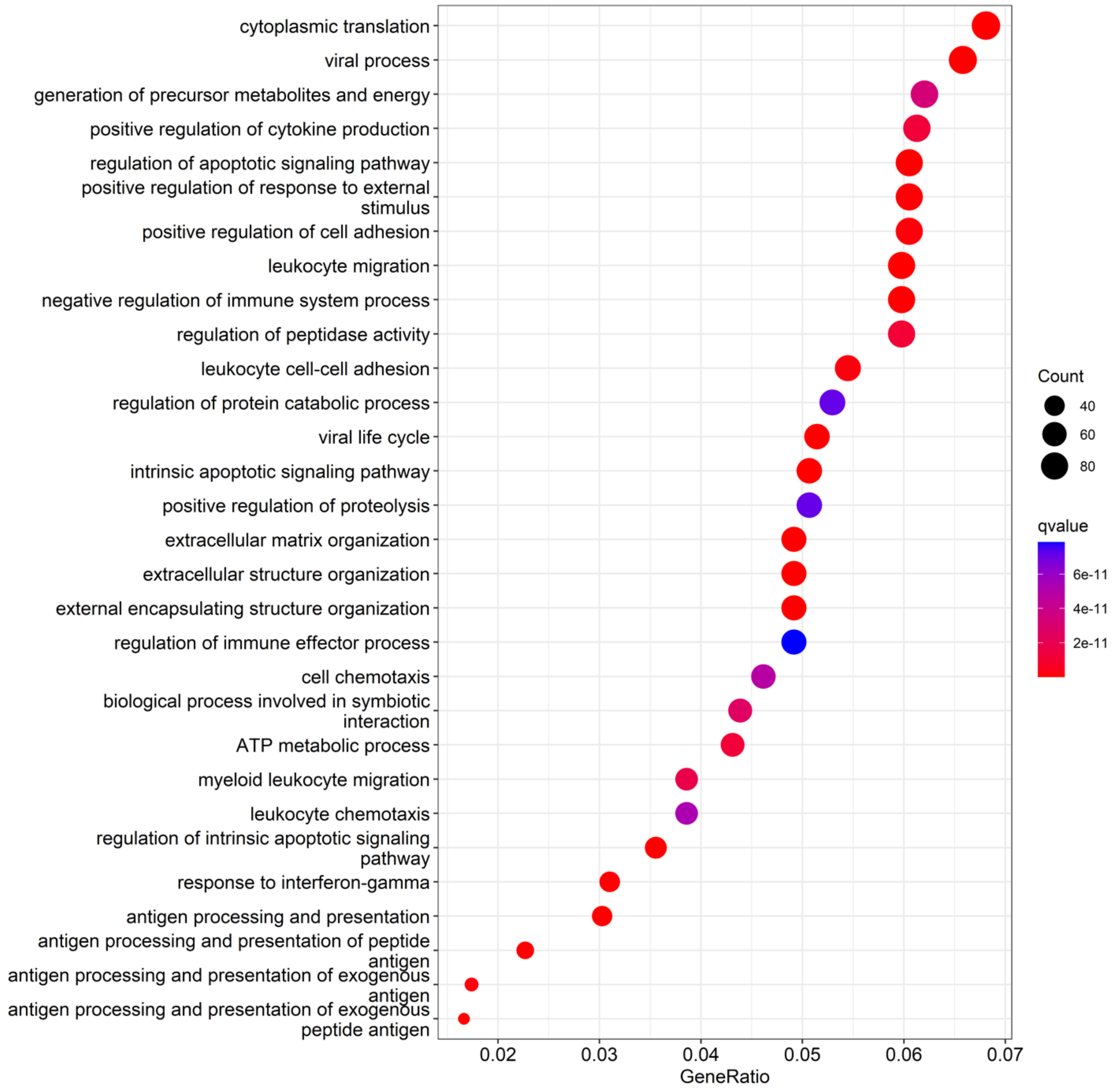


**Supplementary Figure 19: Gene ontology enrichment analysis on patient RA4.**

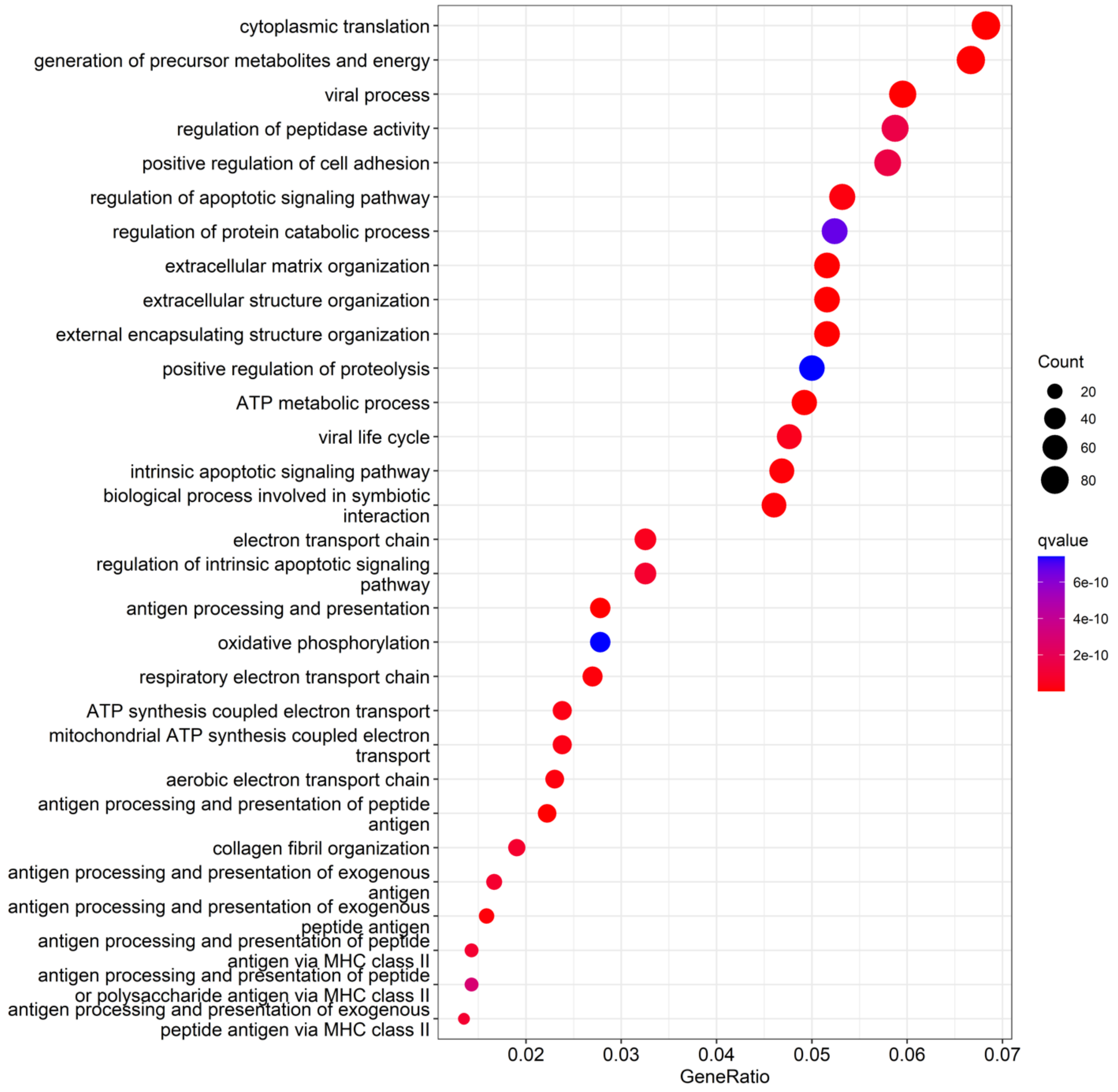


**Supplementary Figure 20: Gene ontology enrichment analysis on patient RA5.**

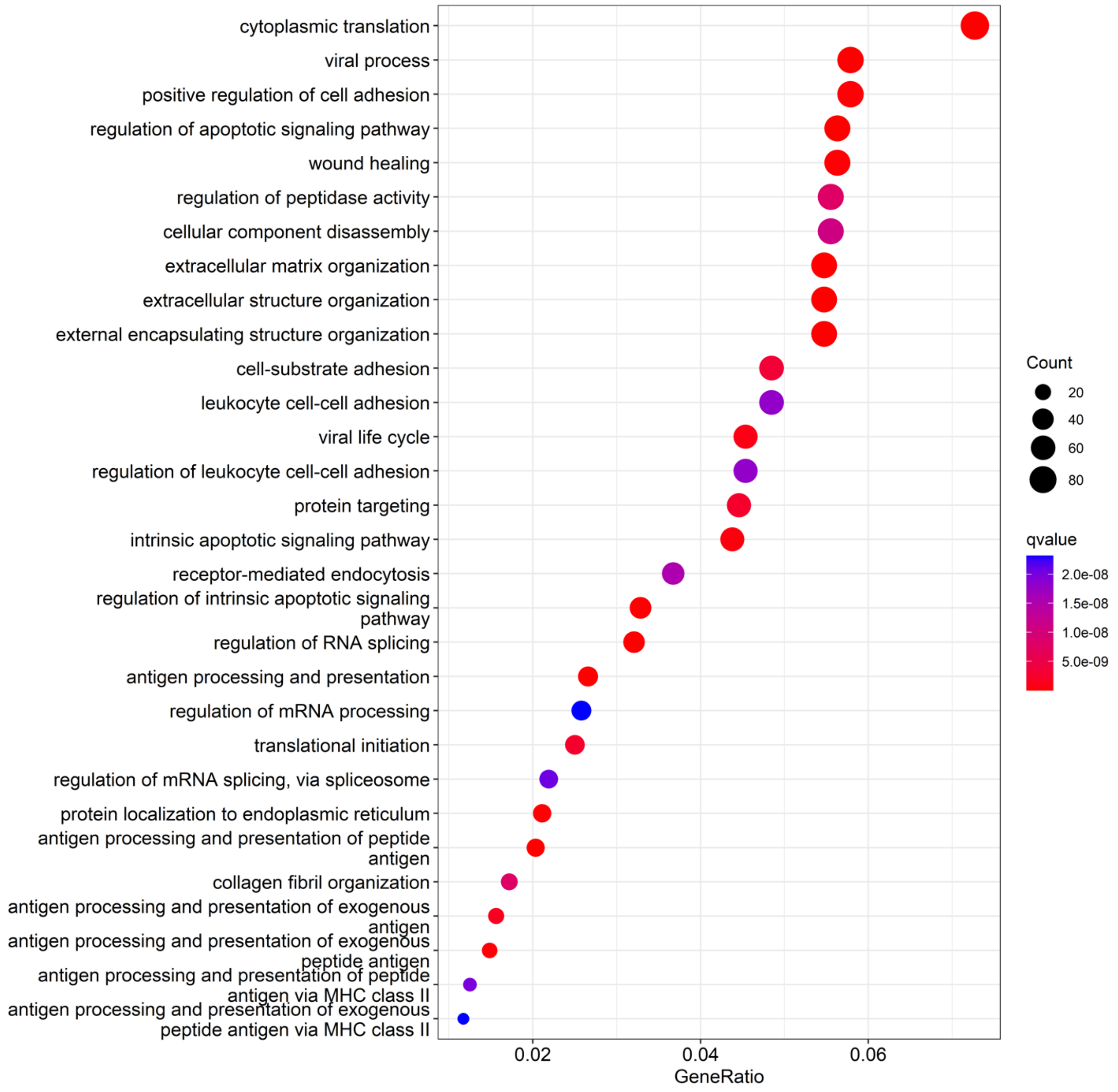


**Supplementary Figure 21: Gene ontology enrichment analysis on patient RA6.**

**Algorithm 1** *BSP* a Big-Small Patch algorithm identifying spatially variable genes in spatial transcriptomics.

**Input:** For spot $i$ , gene expression $\{X_i^{(j)} | 1 \leq i \leq M, 1 \leq j \leq N\}$ for gene $j$, 2D spot coordinates $\{(x_i, y_i)\}$ or 3D spot coordinates $\{(x_i, y_i, z_i)\}$; radius $D_1$ and $D_2$
**Output:** P-value of all genes
Normalize gene expression $X_i^{(j)}$
Normalize coordinates according to $f$ using Eq. (1) or Eq. (2)
**for** gene $j = 1, \ldots, N$ **do**
    **for** spot $i = 1, \ldots, M$ **do**
        Define a small patch $S_i{}'$ with radius $D_1$
        Calculate Local Mean $\widehat{\mathrm{X}}_i^{(j)}$ of small patch $S_i{}'$ using Eq. (3)
    Calculate variance $\sigma_{D_1}^{(j)}$ of Local Mean in all small patches using Eq. (4)
    **end for**
    **for** spot $i = 1, \ldots, M$ **do**
        Define a big patch $S_i{}''$ with radius $D_2$
        Calculate Local Mean $\widehat{\mathrm{X}}_i^{(j)}$ of big patch $S_i{}''$ using with Eq. (3)
    **end for**
    Calculate variance $\sigma_{D_2}^{(j)}$ of Local Mean in all big patches using Eq. (4)
    Calculate $w_j$ as expression normalized factor
    Calculate $r_{D_1 D_2}^{(j)} = w_j \, \sigma_{D_2}^{(j)} / \sigma_{D_1}^{(j)}$ for each gene $j$ as Eq. (5)
Fit a beta distribution of all gene's $\{r_{D_1 D_2}^{(j)}\}$, calculate and adjust P-value for all genes
**end for**

**Supplementary Figure 22: Flowchart of BSP algorithm.**
